# Genetically encoded fluorescent sensors for visualizing polyamine levels, uptake, and distribution

**DOI:** 10.1101/2024.08.21.609037

**Authors:** Ryo Tamura, Jialin Chen, Marijke De Jaeger, Jacqueline F. Morris, David A. Scott, Peter Vangheluwe, Loren L. Looger

**Affiliations:** Department of Neurosciences, University of California, San Diego, La Jolla, CA, USA; Department of Cellular and Molecular Medicine, KU Leuven, Leuven, Belgium; Cancer Metabolism Core, Sanford Burnham Prebys Medical Discovery Institute, La Jolla, CA, USA; Howard Hughes Medical Institute, University of California, San Diego, La Jolla, CA, USA

## Abstract

Polyamines are abundant and physiologically essential biomolecules that play a role in numerous processes, but are disrupted in diseases such as cancer, and cardiovascular and neurological disorders. Despite their importance, measuring free polyamine concentrations and monitoring their metabolism and uptake in cells in real-time remains impossible due to the lack of appropriate biosensors. Here we engineered, characterized, and validated the first genetically encoded biosensors for polyamines, named iPASnFRs. We demonstrate the utility of iPASnFR for detecting polyamine import into mammalian cells, to the cytoplasm, mitochondria, and the nucleus. We demonstrate that these sensors are useful to probe the activity of polyamine transporters and to uncover biochemical pathways underlying the distribution of polyamines amongst organelles. The sensors powered a high-throughput small molecule compound library screen, revealing multiple compounds in different chemical classes that strongly modulate cellular polyamine levels. These sensors will be powerful tools to investigate the complex interplay between polyamine uptake and metabolic pathways, address open questions about their role in health and disease, and enable screening for therapeutic polyamine modulators.

## INTRODUCTION

Polyamines (PAs) are a structurally diverse family of naturally occurring polycationic and aliphatic compounds that play numerous roles in cell biology across all kingdoms of life^1,2^. Putrescine, spermidine, and spermine are in general the most abundant PAs in mammalian cells. PAs are highly positively charged and bind to DNA, RNA, proteins, phospholipids, and sulfated glycans^3–6^, modulating their stability and functions^7–10^. PAs tightly chelate divalent metal ions such as Fe^2+^ and Co^2+^, inhibiting their potent oxidizing ability^11^. Furthermore, the amine groups can act as direct acceptors of free radicals and singlet oxygen^12^, thus serving a secondary antioxidant role^13^. PAs also serve as precursors for post-translational modifications, including hypusination of the translation initiation factor elF5A^14^ and polyamination of tubulin^15^ and many other proteins^16^, regulating their functions. At the cellular level, PAs contribute to DNA repair^17^, maintenance of redox balance^18^, stimulation of autophagy^19^, protein translation induction^20^, regulation of cell morphology^21^, mitochondrial and lysosomal homeostasis^22^, and other processes, ultimately orchestrating the growth, proliferation, differentiation, and survival of various cell types^23–28^.

Cells closely monitor and fine-tune their PA levels by coordinating PA import and export *versus* synthesis and degradation to prevent negative consequences of PA deprivation or toxicity. Ornithine decarboxylase (ODC) and adenosylmethionine decarboxylase (AdoMetDC) represent the rate-limiting steps in PA synthesis, whereas spermidine/spermine N^1^-acetyltransferase 1 (SSAT) and peroxisomal polyamine oxidase (PAOX) catalyze their degradation^29^. The endo-/lysosomal P5B-type transport ATPases ATP13A2-5 have been identified as key contributors to cellular PA uptake by transporting endocytosed polyamines to the cytosol. Only the late endo/lysosomal ATP13A2 and early/recycling endosomal ATP13A3 PA transporters are ubiquitously expressed^30^. Although ATP13A2 and ATP13A3 both contribute to cellular PA uptake^31–36^, ATP13A3 seems to be the principal regulator of PA uptake in mammalian cells, driving faster and more robust PA uptake than ATP13A2, which primarily resides in late endosomal compartments^34^. Also, ATP13A3, but not ATP13A2, is upregulated when PA synthesis is blocked with DL-α-difluoromethyl ornithine (DFMO, a competitive inhibitor of ODC), making ATP13A3 an attractive drug target for PA depletion strategies in cancer^36^.

Dysregulation of PA metabolism and trafficking can be detrimental to human health in multiple aspects. Cancer cells upregulate PA pathways to enable excessive proliferation. Several cancers are associated with altered gene expression of PA-modulating enzymes such as ODC, SSAT, ATP13A3 and ATP13A4, and solute carrier SLC3A2^37–40^. Additionally, mutations in spermine synthase (*SMS*) are causal for the X-linked intellectual disability Snyder-Robinson syndrome, whereas genetic mutations in *ATP13A2* cause Kufor-Rakeb syndrome, a juvenile onset parkinsonism with dementia, and Parkinson’s disease, consistent with PAs’ roles in neurodevelopment and neurodegeneration^41–43^. Mutations in *ATP13A3* have been associated with pulmonary arterial hypertension, a fatal cardiovascular disease^33^. Changes in PA levels in biofluids have been proposed as candidate biomarkers for neurodegeneration and early diagnosis of cancer. To gain insight into the normal physiological roles of PAs and the mechanistic details of disruptions in PA homeostasis in disease, it is critical to develop methods for the real-time, high-resolution, cell type-specific visualization of PAs.

Current methods for PA detection are quite limited, with the state-of-the-art being radioisotopically or fluorescently labeled PA analogs^44–46^. Such probes have proven valuable in quantifying activity of PA transporters, measuring cellular PA uptake, and visualizing PA distribution^34,46^, although radiolabeling has very poor spatial and temporal resolution. Also, being exogenously added, the synthetic probes do not necessarily reflect endogenous PA levels and don’t allow studying the tight interplay between PA uptake and metabolism. Further, PA functionalization by direct conjugation with synthetic dyes alters enzyme and transporter binding and (sub-)cellular uptake, which may complicate interpretation of results^34^.

A wide variety of small molecule dyes that generate fluorescent changes upon binding PAs have been developed^47^. Such dyes can have large and rapid fluorescence responses to PA binding, but they are predicated on supramolecular polymers such as the cucurbit[7]uril (CB[7]) cage. These molecules have poor cellular and tissue availability, are quickly eliminated from cells and blood, and as such provide no path forward for chronic imaging of PA levels in cells or animals. Furthermore, it is impossible to target such molecules to cells or organelles of interest. These probes have only been used in *ex situ* conditions, such as testing on withdrawn blood serum and urine^48^ and stored food for evidence of spoilage^49^. Furthermore, these probes are difficult and expensive to synthesize and are thus used sparingly.

Genetically encoded fluorescent protein-based biosensors are a promising approach to detect diverse analytes of interest^50^, but are not yet available for PAs. Such sensors would offer numerous advantages versus labeled PAs or small molecule dyes: (1) ease of *in situ* and *in vivo* deliverability to measure endogenous PA pools; (2) targetability to cell types of interest and even subcellular organelles; (3) independence from synthetic derivatization, which might perturb pathways; (4) compatibility with long-term and longitudinal imaging, (5) uniform distribution of probe across labeled cells and tissue, and (6) not artificially increasing the PA pool with the labeled probes, thus not disturbing the interplay between PA transport and synthesis. Ideally, PA sensors should respond to physiologically relevant PA concentrations.

Spermidine is generally abundant in mammalian cells and tissues, although concentrations vary widely across cell types, organelles, and conditions, due to heterogeneity in PA biosynthesis, transport, and metabolism. Total PA levels in cells can reach as high as 0.1-30 mM, given that they accumulate due to their membrane impermeance and their association with negatively charged molecules^9,51^. In bovine lymphocytes, total spermidine and spermine concentrations were determined to be 1.3 mM and 1.6 mM, respectively, of which 5-15% and 2-5% represent free (unbound) levels, with most cellular PAs being bound to RNA, DNA, ATP, and phospholipids^52^. The cytosolic concentration of free spermidine and spermine is therefore estimated at 10-100 µM^4,53^. Estimates of PA levels in organelles (nucleus, mitochondria, endosomes, lysosomes, *etc*.) are scarce, although there is evidence of accumulation in each^53^. The effects of PAs on proteins such as K^+^ channels and NMDA receptors manifest at high nanomolar to low micromolar concentrations^54,55^. Thus, PAs exert their influences on cellular processes across a wide concentration range.

Polyamines are ubiquitous in nature and are unsurprisingly consumed by microbes as rich sources of nitrogen^56^. We have previously established a robust pipeline for engineering and optimizing genetically encoded engineered biosensors, using microbial periplasmic binding proteins (PBPs), a class of naturally occurring bacterial biosensors, as sensing modalities. PBPs bind a wide array of environmentally abundant small molecules, including metal ions, carbohydrates, amino acids, peptides, and many more - serving as ideal scaffolds to exploit for biosensor design^50^. Several PA-specific PBPs are known, including PotF and PotD, which exhibit high affinity and selectivity for putrescine and spermidine, respectively^57,58^. Previously, we designed a Förster resonance energy transfer (FRET) polyamine sensor, FLIP-AF1, by fusing CFP at the N-terminus and YFP at the C-terminus of AF1, a PotD homolog from *Agrobacterium tumefaciens* (US20090178149A1). However, FLIP-AF1 has yet to find application due to its limited sensitivity. Here, we present first-in-class single-wavelength PA biosensors (intensiometric polyamine-sensing fluorescent reporters; iPASnFRs) with high signal-to-noise ratio and appropriate affinity for *in situ* deployment. iPASnFRs will be useful for addressing diverse fundamental questions about PA distribution, signaling, and metabolism, and they directly enable screening large libraries of drug candidates for molecules affecting PA handling by cells.

## RESULTS

### Engineering and *in vitro* characterization of genetically encoded polyamine sensors

We have previously shown that PBPs from hyperthermophilic microbes serve as excellent scaffolds for engineering biosensors with high stability and cellular tolerance. We identified multiple PBPs from hyperthermophilic microbes with high sequence identity and homology to *E. coli* PotD (EcPotD hereafter), particularly in the ligand-binding pocket. One of the closest thermophilic homologues was from *Parageobacillus caldoxylosilyticus* (PcPotD hereafter)^59^: PcPotD is 42% identical to EcPotD (**Supp. Fig. 1**). Superimposition of the PcPotD structure predicted by AlphaFold2^60^ and the crystal structure of EcPotD (PDB: 1POT)^61^ shows that the spermidine-binding residues of EcPotD are generally conserved in PcPotD, suggesting that PcPotD likely binds spermidine and/or related polyamines (**Supp. Fig. 1**).

To convert PcPotD into a fluorescent reporter, we inserted circularly permuted superfolder Venus (cpsfVenus)^62^ into PcPotD at multiple locations: G47, E66, G68, E284, D291, all of which lie in loops in which the fluorescent protein is plausibly allosterically modulated by the open-to-closed “Venus flytrap” mode of ligand binding-induced PBP conformational change (**Supp. Fig. 2A**). All of these prototype sensors expressed in *E. coli* and were titrated with spermidine. The prototype sensors with insertions at E66 and E284 responded (**Supp. Fig. 2B**), of which the most promising was PcPotD-cpsfVenus-E284, with a maximum fluorescence response (ΔF/F_0_)_max_ of ∼0.6 and a K_d_ of 0.6 µM (**Supp. Fig. 2C**). We therefore selected it as a starting variant for sensor engineering. To optimize fluorescence intensity increase and sensitivity for spermidine, we screened libraries of PcPotD-cpsfVenus mutants randomized at the two linkers joining PcPotD and sfVenus, and surrounding residues. Screening ∼20,000 variants produced a clone with (ΔF/F_0_)_max_ of ∼11 and a K_d_ of 5.5 µM for spermidine; this clone had two altered linkers (-SK- and -YH-) and 2 additional mutations: Lys19Arg and Asn287Arg (**Fig. 1A and Supp. Fig. 1A**). We termed this variant iPASnFR-H, due to this variant’s relatively high affinity for spermidine.

**Figure 1:**
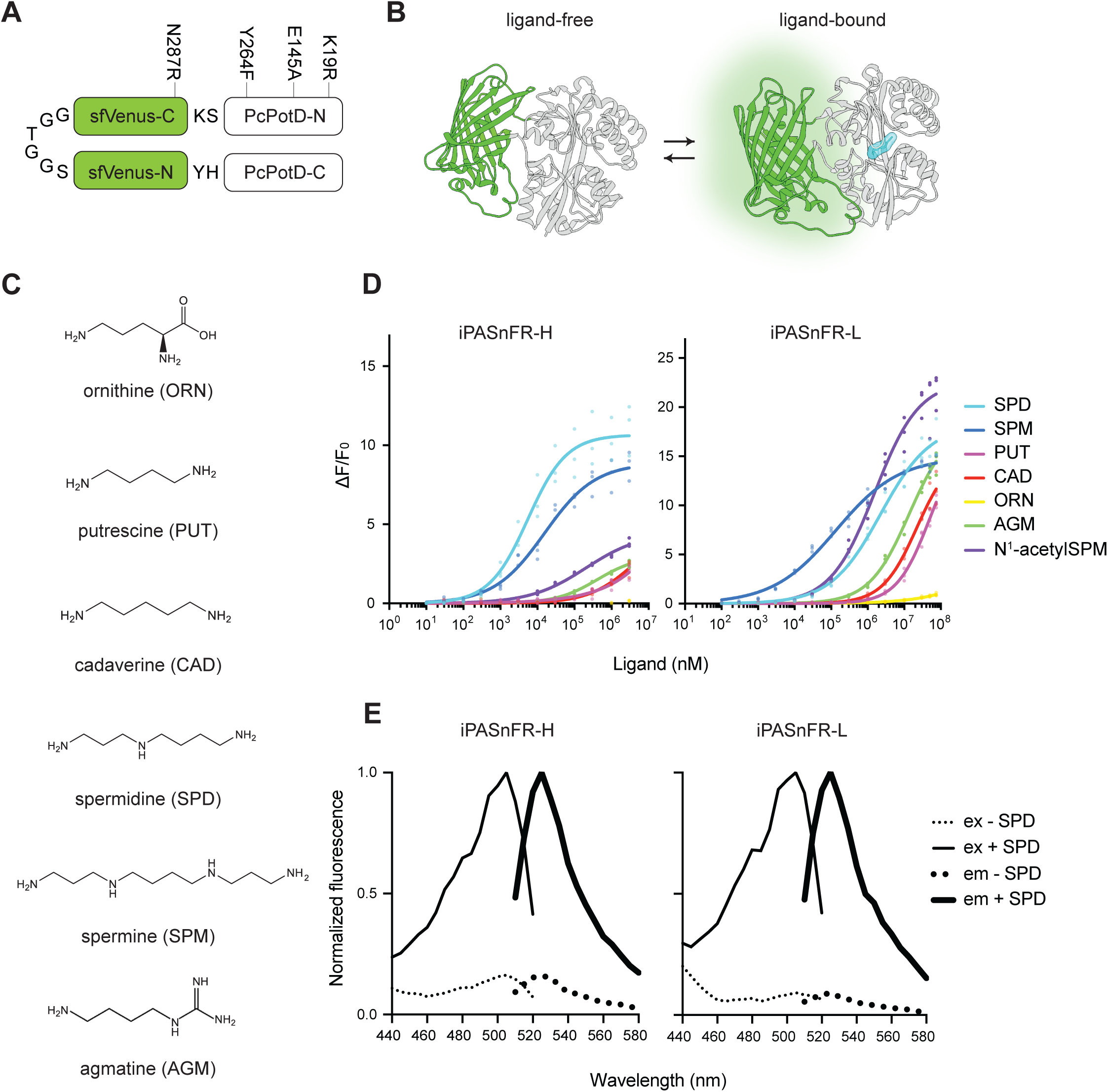
Engineering and *in vitro* characterization of genetically encoded polyamine sensors. (**A**) Primary structure of iPASnFR-H/L. Denoted are mutations in PcPotD (white) or sfVenus (light green) discovered during sensor optimization. N: N-terminus; C: C-terminus. (**B**) Mechanism of iPASnFR-H/L. Upon ligand (cyan) binding, the binding protein undergoes conformational changes that are transduced to cpsfVenus, causing fluorescence increase. Structural models of ligand-free (left) and ligand-bound (right) sensors are shown. (**C**) Chemical structures of diamines: putrescine, and cadaverine; a triamine: spermidine; a tetraamine: spermine; and the precursor molecules ornithine and agmatine. (**D**) Titration of purified iPASnFR-H/L proteins with polyamines shown in (**C**) (*n* = 3). ΔF/F_0_: (F – F_0_) / F_0_. (**E**) Excitation and emission spectra of iPASnFR-H/L in the presence (bold lines) or absence (thin lines) of 10 mM spermidine.

To fully understand the biology of PAs and their diverse roles in signaling and homeostasis, it is desirable to have a series of biosensors spanning as much of the physiological concentration range as possible. To make a sensor useful for sensing at high concentrations, we generated a low-affinity version of iPASnFR by substituting glutamate 145, modeled to bind to a spermidine amine moiety, to alanine (Glu145Ala), and Tyr264Phe, which lies adjacent to the linker-cpVenus junction (**Fig. 1A and Supp. Fig. 1A**). The resulting variant displays a (ΔF/F_0_)_max_ of ∼18 and a K_d_ of 2.3 mM for spermidine (**Fig. 1D**), yielding iPASnFR-L (low affinity). As expected, iPASnFR-H/L showed excitation/emission maxima similar to those of their parent mVenus^63^ (**Fig. 1E**). Together, iPASnFR-H and iPASnFR-L span most of the range of PA levels expected to be relevant in cell biology. However, the precise levels of PAs across cells, organelles, and cell states are essentially unknown at this point.

Diverse polyamines play distinct roles in the cell. Therefore, it is vital to characterize the ligand-binding specificity of iPASnFR-H/L. Titration of iPASnFR-H/L with different polyamines (spermidine, spermine, putrescine, and cadaverine; also the analogue agmatine) and their amino acid precursor ornithine (**Fig. 1C**) showed that iPASnFR-H/L preferentially respond to spermidine and spermine (**Fig. 1D**), consistent with the reported preference of EcPotD^57,58^: iPASnFR-H has a (ΔF/F_0_)_max_ of ∼8.9 and a K_d_ of 15 µM and iPASnFR-L a (ΔF/F_0_)_max_ of ∼15 and a K_d_ of 0.13 mM for spermine, comparable to their responses towards spermidine. Additionally, iPASnFR-L has a (ΔF/F_0_)_max_ of ∼23 and a K_d_ of 1.6 mM for N^1^-acetylspermine, although iPASnFR-H did not saturate up to 10 mM, suggesting altered binding modes of the two sensors, perhaps mediated through repulsion between Glu145 in iPASnFR-H and the acetyl moiety. By contrast, iPASnFR-H/L have markedly lower sensitivity for putrescine (iPASnFR-H: ΔF/F_0_ of ∼1.3; iPASnFR-L: ΔF/F_0_ of ∼0.47 at 1 mM putrescine) and agmatine (iPASnFR-H: ΔF/F_0_ of ∼3.0; iPASnFR-L: ΔF/F_0_ of ∼1.9 at 1 mM agmatine) and are non-responsive to ornithine and cadaverine in their physiological ranges. Thus, iPASnFR-H/L will mostly report spermidine, spermine, N^1^-acetylspermine, and to a lesser extent putrescine and agmatine. *In vitro* pH titration with purified proteins shows that iPASnFR-H/L decrease in saturated brightness, with a p*K_a_* of ∼6.8 and ∼6.5, respectively. The ΔF/F_0_ of iPASnFR-L, however, remains ∼0.40 at pH 5.0 and ∼1.7 at pH 5.5 (**Supp. Fig. 3**), indicating its potential utility in acidic compartments like endosomes and lysosomes.

### iPASnFR detects drug-induced changes in cytosolic PA levels and demonstrates synergistic effects of inhibiting PA synthesis and uptake

To demonstrate the utility of iPASnFRs in sensing intracellular PAs, we created mammalian expression plasmids encoding iPASnFRs fused to the bright, photostable, monomeric red fluorescent protein mScarlet3^64^ (**Fig. 2A**), as a reference fluorophore. This allows us to standardize sensor fluorescence relative to its expression in individual cells and compartments by computing the ratio of emission between iPASnFR and mScarlet3 (‘normalized fluorescence’). Critically, we confirmed that PAs did not affect mScarlet3 fluorescence (discussed in **Fig. 3D** and **Supp. Fig. 4A-B**); thus, it functions as an inert signal of sensor abundance in a given field-of-view. When transiently expressed in the cytoplasm of HEK293, iPASnFR-H displays a ∼1.8-fold higher normalized fluorescence than iPASnFR-L, in agreement with the affinity difference of iPASnFR-H/L for PAs determined *in vitro* (**Fig. 2B**).

**Figure 2:**
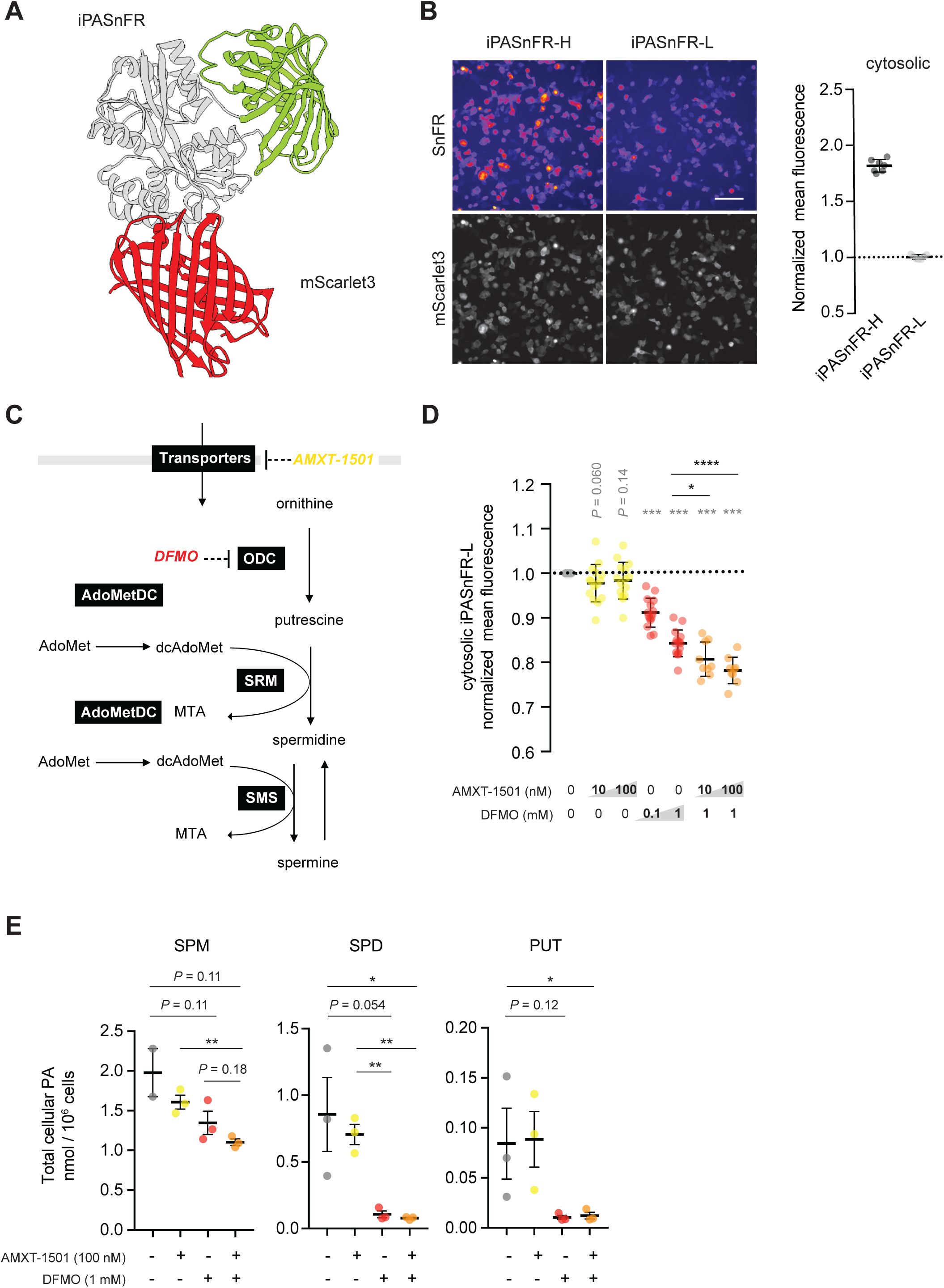
iPASnFR detects drug-induced changes in cytosolic PA levels and demonstrates synergistic effects of inhibiting PA synthesis and uptake. (**A**) Structural cartoon of iPASnFR-mScarlet3 (gray: binding protein; light green: cpsfVenus; red: mScarlet3). (**B**) Cytosolically expressed iPASnFR-H/L in HEK293 cells. Scale bar, 100 µm. The ratio of iPASnFR to mScarlet3 in individual cells was quantified and averaged in each replicate, covering ∼500 cells. Seven replicates from 2 independent experiments are shown. The mean ratios were normalized to the average of those of iPASnFR-L. (**C**) Schematic of PA biosynthetic pathway. ODC: ornithine decarboxylase; SRM: spermidine synthase; SMS: spermine synthase; AdoMetDC: S-adenosylmethionine decarboxylase; dcAdoMet: decarboxylated S-adenosylmethionine; MTA: methylthioadenosine. AMXT-1501 and DFMO are PA analogs that competitively inhibit PA transporters and ODC, respectively. (**D**) Measurement of mean iPASnFR-L:mScarlet3 fluorescence ratio in HEK293 cells treated with AMXT-1501 and DFMO either individually or in combination. A total of 9-15 replicates from 2-3 independent experiments is shown for each condition. We used the indicated drug concentrations, to which iPASnFR-L is not sensitive (DFMO, ΔF/F_0_: −0.069 ± 0.020 at 1 mM; AMXT-1501, ΔF/F_0_: 0.049 ± 0.0080 at 100 nM) (**Supp. Fig. 6**). The mean fluorescence ratios were normalized to control (no drug addition) in each experiment. Single-sample *t*-tests were performed to compare mean fluorescence ratios of drug-treated groups to non-treated (gray). Comparisons between groups were made using 2-tailed unpaired *t*-tests (black). (**E**) Metabolomics analysis of total cellular PAs (SPM, SPD, and PUT) in HEK293 cells treated with DFMO and AMXT-1501 either alone or in combination. Two-tailed unpaired *t*-tests were performed for pairwise analyses. When *P* > 0.05, individual *P*-values were shown above comparison bars. No significant differences were observed unless noted.

**Figure 3:**
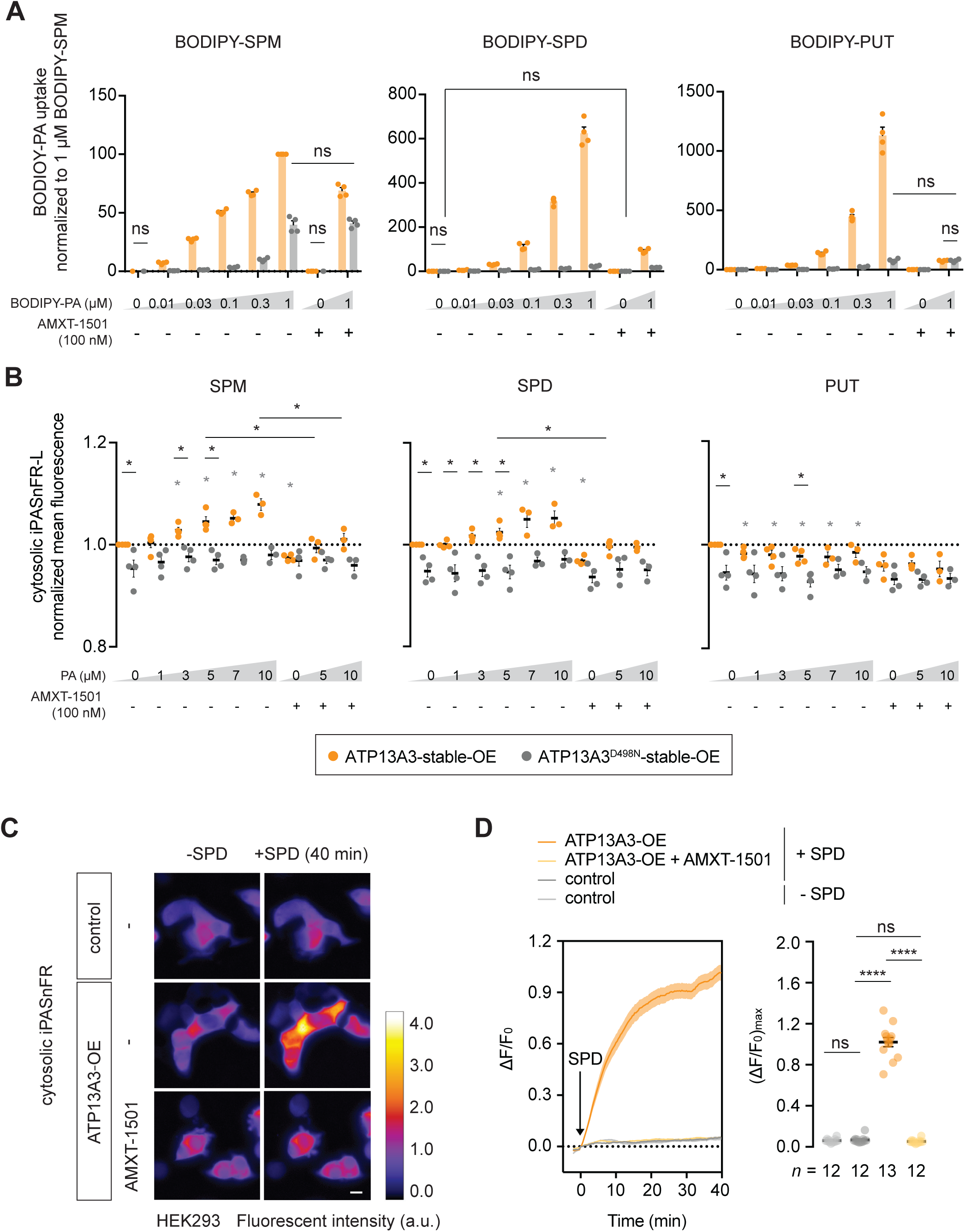
iPASnFR-L responds to ATP13A3-mediated cytosolic PA uptake. (**A**) Mean fluorescence intensities (MFIs) of ATP13A3-stable-OE and ATP13A3^D498N^-stable-OE HEK293T cells after dose-dependent supplementation of BODIPY-PAs for 0.5 h. Uptake experiments were also performed with 100 nM AMXT-1501 at 0 and 1 μM BODIPY-PAs. Results were normalized to ATP13A3-stable-OE cells supplemented with 1 μM BODIPY-SPM. (**B**) ATP13A3-stable-OE and ATP13A3^D498N^-stable-OE cells with transient expression of cytosolic iPASnFR-L-mScarlet3 were supplemented with indicated amounts of unmodified SPM, SPD and PUT for 0.5 h. Uptake experiments were also performed with 100 nM AMXT-1501 at 0, 5 and 10 μM PA. MFIs of iPASnFR-L were divided by mScarlet3 intensities for expression correction, then normalized to non-treated ATP13A3-stable-OE cells. In (A), all pairs of comparisons between ATP13A3-stable-OE and ATP13A3^D498N^-stable-OE cell lines at the same BODIPY-PA concentrations were significantly different unless noted. (**C**) Confocal images of HEK293 cells expressing iPASnFR-L (with or without ATP13A3 co-expression), with or without AMXT-1501 treatment. Left: pre-spermidine addition; right: post-spermidine addition at 40 min. Scale bar, 10 µm. (**D**) Live tracking of iPASnFR-L signal (mean ± s.e.m.) over time upon spermidine addition at 0 min, corresponding to (B). *n* = 12-13 cells from 2 independent experiments. Two-tailed Mann–Whitney *U* tests were performed for pairwise analyses.

Therefore, we reasoned that iPASnFR-L would prove more useful than iPASnFR-H for detecting changes in free cellular PA levels, given its response to spermidine and spermine in the physiological range (10 µM - 10 mM) (**Fig. 1D**). To validate the sensitivity of iPASnFR-L in cultured cells, we challenged transfected HEK293 cells with three PA pathway-modifying compounds, either individually or in combination (**Supp. Figs. 5-6**): AMXT-1501, a PA transport inhibitor preventing cellular PA uptake, and DFMO, a competitive inhibitor of ODC (**Fig. 2C**). Individual drug treatments showed that AMXT-1501 alone does not substantially decrease the normalized fluorescence (most likely reflecting combined free cytosolic spermidine and spermine concentrations) within the detection range of iPASnFR-L, whereas 1 mM DFMO does so by ∼15%, indicating that HEK293 cell may rely more on PA synthesis than uptake under basal conditions. Moreover, 1 mM DFMO makes HEK293 cells more reliant on PA uptake to maintain cytosolic PA levels, since AMXT-1501 lowers the free PA concentration further during DFMO treatment by ∼6% in iPASnFR-L signal (**Fig. 2D**). This provides evidence that inhibition of biosynthesis and cellular uptake act synergistically on the free cytosolic PA levels, in support of PA depletion strategies with DFMO and AMXT-1501 for cancer therapy^37^. In fact, previous studies in neuroblastoma cancer cells^36^ suggest that DFMO treatment induces upregulation of ATP13A3, and that cellular ATP13A3 activity is blocked by AMXT-1501, corroborating this notion - and that cells respond to PA biosynthesis blockade by upregulating import.

To validate the observations from iPASnFR-L in HEK293 cells with an independent approach, we performed metabolomics analysis. Detection of total cellular putrescine, spermidine, and spermine with gas chromatography-mass spectrometry was performed using HEK293 treated with DFMO and AMXT-1501, either individually or in combination. We found that DFMO-only treatment substantially decreased total PA amount (total spermine: −32%; spermidine: −85%; putrescine −88%) compared to AMXT-1501-only treatment (total spermine: −19%; spermidine: −22%; putrescine −5%) (**Fig. 2E**), supporting the largest decrease we observed in iPASnFR-L signal with DFMO among other individual drug treatments (**Fig. 2D**). AMXT-1501 treatment in the presence of DFMO further decreased total spermine content (**Fig. 2E**) by ∼25%, consistent with the iPASnFR-L observations with DFMO alone *versus* DFMO and AMXT-1501 (**Fig. 2D**). We observed larger decreases in the total spermine and spermidine content *via* metabolomics than with iPASnFR-L. This may reflect 1) the inherent difference in the detection methods, in which iPASnFR-L exclusively detects cytosolic, free spermine and spermidine, whereas mass spectrometry with total cell lysates detects total PA contents in all cellular compartments, 2) the large buffering capacity of intracellular RNA, DNA, ATP, and lipids for PA binding, 3) the response of iPASnFR-L towards multiple PA species at different affinities, including acetylated spermine, which was not detected by this metabolomics analysis (**Fig. 1D**). Nonetheless, these data confirm that iPASnFR-L is faithfully reporting PA fluctuations in the cell. The metabolomics analysis also revealed that the abundance of putrescine in untreated HEK293 cells was ∼0.04% relative to spermine and ∼0.1% relative to spermidine, suggesting that iPASnFR-L would essentially report just spermine, spermidine and their acetylated derivatives under physiological conditions.

Altogether, these results demonstrate the utility of iPASnFR-L in capturing drug-induced changes in endogenous free PA levels in cultured cells. Furthermore, this opens up an opportunity for large-scale drug discovery targeting endogenous PA pathways.

### iPASnFR-L responds to ATP13A3-mediated cytosolic PA uptake

The ATP13A3 transporter has emerged as the major component of the mammalian PA transport system. Elevated ATP13A3 activity evokes fast and robust uptake of putrescine, spermidine and spermine, regulating the total cellular PA content^32–36^. We therefore validated the sensitivity and specificity of the iPASnFR-L sensor in an ATP13A3 gain-of-function model supplemented with putrescine, spermidine or spermine.

Given that variation in transporter expression via transient transfection could result in large differences in PA flux, we first established a stably integrated HEK293T line expressing wild-type ATP13A3 (ATP13A3-stable-OE), as well as a stable line of the transport-dead ATP13A3 mutant D498N (ATP13A3^D498N^-stable-OE) as a negative control. We first verified expression of functional ATP13A3 by measuring cellular uptake of fluorescent substrates BODIPY-labeled putrescine (BODIPY-PUT), spermidine (BODIPY-SPD), and spermine (BODIPY-SPM), which are substrates of the P5B-type PA transporters^34^. As expected, the wild-type transporter mediated uptake of BODIPY-PUT, BODIPY-SPD, and to a lesser extent BODIPY-SPM in a dose-dependent manner (**Fig. 3A**), as previously observed in other cell models with modified ATP13A3 expression^33,34,65^. Conversely, the dead transporter showed substantially lower BODIPY-PA uptake. Further, the rise in fluorescence in ATP13A3-stable-OE cells was inhibited by AMXT-1501 (**Fig. 3A**), as observed before^36^, confirming that the flux is indeed through ATP13 transporters.

We then transiently transfected iPASnFR-L into these two HEK293T cell lines, and imaged biosensor fluorescence responses to unmodified PAs instead of fluorescent PA analogues. The basal normalized fluorescence of iPASnFR-L was higher in ATP13A3-stable-OE cells versus ATP13A3^D498N^-stable-OE cells (**Fig. 3B**), consistent with influx occurring through wild-type ATP13A3, but not ATP13A3^D498N^. Moreover, the free PA concentration gradually and dose-dependently increased in ATP13A3-stable-OE cells with spermidine and spermine, but not putrescine (**Fig. 3B**). This selective response to spermine and spermidine validates the above results about sensor selectivity. No sensor response is observed in ATP13A3^D498N^-stable-OE cells, and the response in ATP13A3-stable-OE cells can be blocked with AMXT-1501 (**Fig. 3A**), demonstrating that the higher iPASnFR-L signal is a direct consequence of ATP13A3-mediated PA transport into the cytosol. Thus, iPASnFR-L sensor signal replicates observations from fluorescently labeled substrates.

To investigate the kinetics of cytosolic PA uptake through ATP13A3, we performed live-cell imaging of cytoplasmic iPASnFR-L-mScarlet3 in HEK293 cells transiently overexpressing wild-type ATP13A3 *versus* unmodified cells. Supplementation of external spermidine to control HEK293 cells did not significantly change cytosolic iPASnFR-L signals over the course of 40 min (**Fig. 3C-D**), further confirming that under basal conditions HEK293 control cells mainly rely on PA synthesis rather than PA uptake (**Fig. 3D**). Conversely, transient expression of wild-type ATP13A3 promoted cytosolic spermidine uptake with a t_1/2_ of 6 min, demonstrating fast cytosolic PA uptake mediated by ATP13A3 that reaches a maximal response in 20 min (**Fig. 3D**). This is a similar timeline as that found for ATP13A3-mediated BODIPY-PA uptake^34^, indicating that this time course genuinely represents PA transport into the cytosol. The ATP13A3-dependent spermidine uptake was again blocked by pre-treating cells with AMXT-1501 (**Fig. 3D**). Additionally, we confirmed that spermidine supplementation exclusively affects the iPASnFR-L signal, but not that of fused mScarlet3, confirming the inertness of mScarlet3 as a PA-insensitive reference fluorophore in the iPASnFR-L-mScarlet3 fusion (**Supp. Fig. 4A-B**).

Altogether, these data suggest that the genetically encoded iPASnFR-L sensor is useful for visualizing real-time cytosolic PA uptake through transporter activity, and that results conform with those using other methods.

### ATP13A2 and ATP13A3 play distinct roles in subcellular flux of exogenous PAs

Both ATP13A2 and ATP13A3 contribute to cellular PA uptake and are ubiquitously expressed, suggesting possibly redundant roles, although the transporters are genetically associated with distinct disorders. We directly tested whether the transporters may differentially affect PA homeostasis by using iPASnFR-L to assess their relative effects on cellular PA uptake kinetics, free cytosolic PA levels, and subcellular PA distribution. Importantly, the iPASnFR-L-mScarlet3 fusion constructs correctly trafficked to the nucleus (by adding a triple nuclear localization signal) and the mitochondrial lumen (by adding a Cox8-derived targeting peptide) (**Fig. 4A**). To validate the use of iPASnFR-L in these subcellular locations, we quantified the emission ratio between iPASnFR-L and mScarlet3 (**Supp. Fig. 7A**) and compared the relative abundance of free PAs in the cytosol, nucleus, and mitochondria. We found that iPASnFR-L:mScarlet3 ratios are highest in mitochondria (∼2.7-fold), followed by nucleus (∼1.2-fold), relative to cytosol (**Supp. Fig. 7B**). These data agree with the reported mitochondrial accumulation of PAs in mitochondria^22^. Total mitochondrial PA concentration is estimated at ∼2 mM (spermidine: ∼0.62 mM; spermine: ∼1.6 mM) in rat liver^66^, comparable to the total intracellular PA content estimated in bovine lymphocytes (spermidine: 1.3 mM; spermine: 1.6 mM)^4,53^. Our observation with iPASnFR-L suggests that free PA levels in mitochondria are higher than those in the cytosol or nucleus.

**Figure 4:**
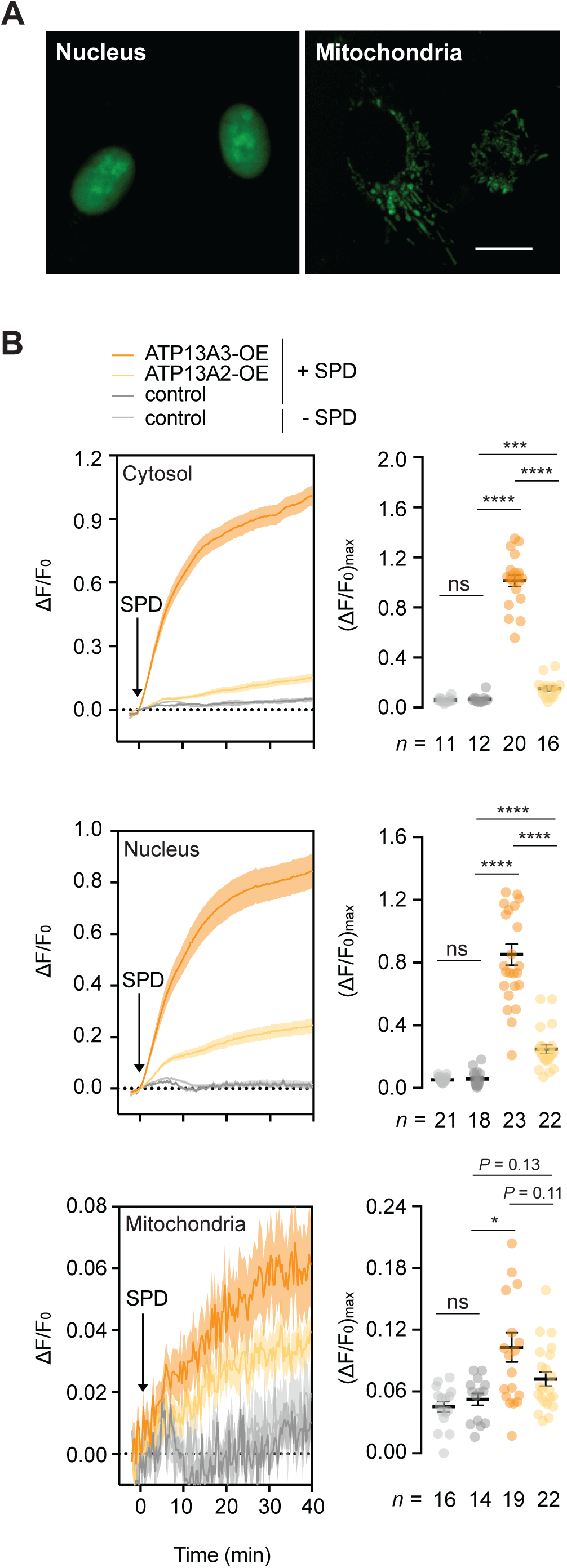
ATP13A2 and ATP13A3 play distinct roles in subcellular flux of exogenous PAs. (**A**) Confocal images of U2OS cells expressing iPASnFR-L in the nucleus or mitochondria. Scale bar, 20 µm. (**B**) Live tracking of iPASnFR-L signal in cytosol, nucleus, or mitochondria over time (mean ± s.e.m.) upon spermidine addition at 0 min with ATP13A2/ATP13A3 co-expression in HEK293 cells. *n* = 11-23 cells from 2-3 independent experiments per condition. Two-tailed Mann–Whitney *U* tests were performed for pairwise comparison. When *P* > 0.05, individual *P* values are shown above comparison bars.

Next, we examined the relative effect of ATP13A2 or ATP13A3 transient expression on the free cytosolic, nuclear and mitochondrial PA levels in HEK293 cells co-expressing iPASnFR-L. Subsequently, spermidine was externally added, and transfected cells were imaged by live fluorescence microscopy in serum-free media to prevent PA degradation from extracellular PA oxidase (present in media). In untransfected HEK293 cells (denoted control), exogenous supplementation of 10 µM spermidine did not significantly increase its concentration in the nucleus and mitochondria as in the cytosol (**Fig. 4B**). Conversely, transient expression of ATP13A3 significantly increases maximum spermidine with a (ΔF/F_0_)_max_ of up to ∼0.85 (nuclei) and ∼0.10 (mitochondria), respectively, compared to untransfected control, suggesting that ATP13A3 mediates nuclear and mitochondrial spermidine influx. In comparison, ATP13A2 expression yields a spermidine influx of only ∼15% compared to ATP13A3 in the cytosol, corroborating that ATP13A3 is a major mediator of cytosolic spermidine influx^32–36^. In the nucleus, however, ATP13A2 expression yields a maximum spermidine uptake of ∼24% relative to ATP13A3, and in mitochondria, ∼40% relative to ATP13A3, indicating that ATP13A2 may exert a more organelle-specific function than ATP13A3 in PA distribution. Indeed, we have previously shown that ATP13A2 redistributes PAs to mitochondria using fluorescently labeled spermine^22^, in agreement with our observations here with iPASnFR-L. The relatively small average (ΔF/F_0_)_max_ of iPASnFR-L in mitochondria (ATP13A3: ∼0.097; ATP13A2: ∼0.075) *versus* those in cytosol (ATP13A3: ∼1.0; ATP13A2: ∼0.15) and nuclei (ATP13A3: ∼0.70; ATP13A2: ∼0.27) can be in part attributed to the high basal free mitochondrial PA level (**Supp. Fig. 7B**), which would decrease the dynamic range of sensor response.

Altogether, these data suggest that iPASnFR-L is useful for visualizing real-time PA dynamics in distinct cellular compartments, which will allow us to ask, for example, how different molecular components affect intracellular PA distribution.

### Small compound screen reveals upstream signaling programs of polyamine metabolism

Cancer cells are known to alter intracellular PA levels to maintain proliferation and survival, in part by altering the transcriptional profile of PA-related enzymes and transporters^36,67,68^. However, the molecular constituents upstream of such reprogramming are only beginning to be deciphered^69–74^. To identify signaling networks involved in PA metabolism that could potentially be targeted for therapeutic development, we screened a commercially available compound library of 1,268 molecules associated with key metabolic pathways and known targets often dysregulated in cancer. To achieve this, we first transiently expressed our fully validated cytosolic iPASnFR-L in HEK293 cells, seeded the cells in 96-well format (compatible with high-throughput imaging), administered small molecule compounds 24 h post-transfection, and imaged them at 24, 48, 72 h post-treatment, followed by identification of compounds that either significantly increased or decreased basal PA levels based on the iPASnFR-L-to-mScarlet3 emission ratio (**Fig. 5A**). A total of 393 compounds produced significant changes in PA levels (217 at 24 h, 132 at 48 h, 130 at 72 h) with *P* < 0.05 and absolute log fold change > 0.05 (**Fig. 5B**), implicated in multiple cellular pathways. Additionally, there was a general trend that large decreases in PA levels peaked at 48 h, whereas large PA increases more gradually manifested at 72 h (**Fig. 5B**), suggesting that PA levels in HEK293 cells are more likely to be resilient to upregulation than downregulation. This observation was further confirmed by time-series analyses of the fold change of the top 25 compounds that either increased or decreased PA levels by the largest margin at 72 h (**Fig. 3C**).

**Figure 5:**
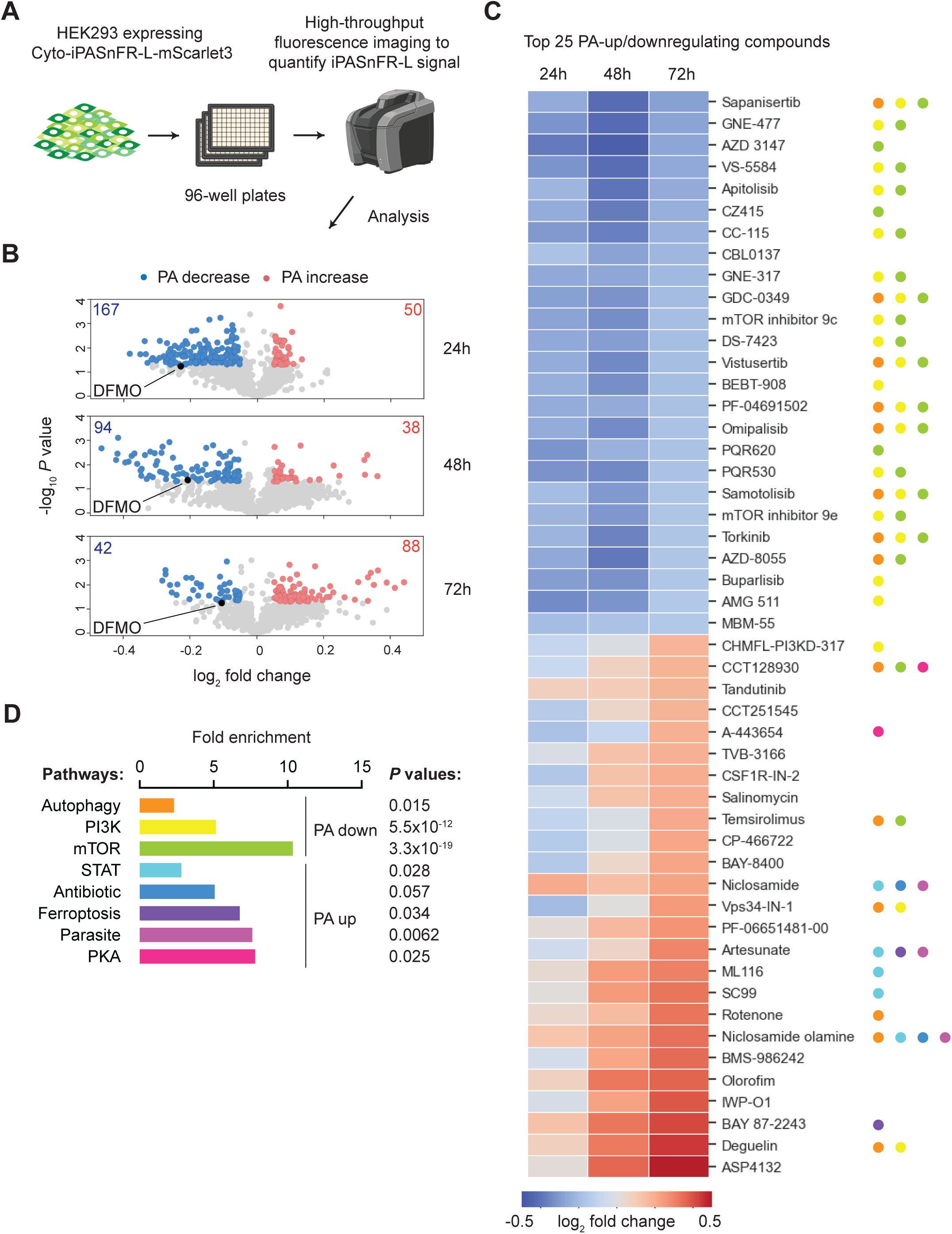
Small compound screen reveals upstream signaling programs of polyamine metabolism. (**A**) Design of compound screen. (**B**) Volcano plots of iPASnFR-L response to individual compounds at 3 time points (24, 48, 72 h). Numbers and individual data points of compounds that significantly (*P* < 0.05) increased (red) or decreased (blue) iPASnFR-L signal (absolute log fold change > 0.05) are indicated. Control samples treated with 1 mM DFMO are shown in black for reference. (**C**) Heatmap showing fold changes of the top 25 compounds either increasing or decreasing iPASnFR-L signal at 72 h. (**D**) Fold enrichment of pathway classes with which the top 25 compounds have been previously associated. Each compound in (**C**) is marked with hues corresponding to those in (**D**) according to their related pathways.

A subsequent pathway enrichment analysis based on these top 50 compounds revealed that inhibitors of the PI3K/AKT/mTOR (phosphoinositide 3-kinase/protein kinase B/mechanistic target of rapamycin) pathway were significantly associated with PA downregulation (mTOR, 10-fold enrichment over expected value, Fisher exact test *P* value: 3.28E-19; PI3K, 5-fold enrichment over expected value, Fisher exact test *P* value: 5.56E-12). Inhibitors of autophagy, which is regulated by mTOR signaling, were also associated with PA downregulation (2-fold enrichment over expected value, Fisher exact test *P* value: 0.015). Indeed, both PI3K and mTOR have been implicated in altered PA metabolism: PI3K activation causes spermidine production, promoting cell survival^75^. Furthermore, the PTEN/PI3K/mTORC1 pathway sustains tumor growth and proliferation by directly stabilizing AdoMetDC and maintaining PA biosynthesis^70^. Overall, the substantial enrichment of the PI3K/AKT/mTOR pathway in our screen results further underscores these previous findings. Additionally, transcriptome analyses based on Gene Expression Omnibus (GEO) database revealed that a number of these polyamine-decreasing compounds decrease expression of key polyamine-synthesizing genes and transporters in parallel with increased expression of polyamine-catabolizing genes. These data suggest that change in PA levels by the identified compounds occur in part through transcriptional regulation downstream of the PI3K/AKT/mTOR pathway (**Supp. Fig. 8**), although further validation is required to pinpoint the precise mechanisms of PA downregulation mediated by each compound.

Conversely, targets significantly enriched in the top 25 PA-increasing compounds were more diverse: Inhibitors targeting the JAK/STAT pathway, key for immune cell responses to various cytokines (STAT, 3-fold enrichment above the expected value, Fisher exact test *P* value: 0.028)^76^, were significantly enriched, revealing the JAK/STAT pathway as a novel upstream signaling cascade affecting resting PA levels. Additionally, antimalarial (artesunate) and anthelminthic/antibiotic (niclosamide and its derivative niclosamide olamine) compounds were significantly associated with PA upregulation (antiparasitic, 8-fold enrichment above the expected value, Fisher exact test *P* value: 0.0062; antibiotic, 5-fold enrichment above the expected value, Fisher exact test *P* value: 0.057). PA biosynthesis is a validated drug target to counter bacterial, parasitic, and helminthic infections^77^, indicating the relevance of PA manipulation as a broadly conserved anti-pathogen strategy - it is not clear why these anti-pathogen compounds were associated with PA upregulation rather than downregulation. ASP4132, the top PA-increasing compound, is an activator of the AMPK pathway: AMPK activation has previously been shown to promote PA biosynthesis by upregulating ODC translation and thus increasing PA levels^69^, further verifying our observations. However, AMPK also serves as an antagonist of the mTOR pathway^78^, highlighting the complexities of the PA metabolic network.

Overall, these results demonstrate that 1) iPASnFR-L is useful in establishing a high-throughput cell-based platform to conduct large-scale compound screens, with multiple drug classes showing large and significant increases or decreases in resting PA levels, and that 2) such screens will allow us to predict how cells may shift the PA metabolic landscape at the system level through signaling in response to external perturbation.

## DISCUSSION

We have created robust genetically encoded biosensors for detecting polyamines (PAs), particularly spermine and spermidine, and to a lesser extent putrescine. Our sensors have excellent binding-dependent fluorescence increase, and high specificity for polyamines over other molecules. We have rigorously characterized the sensors in a variety of applications in cell culture, showing that the sensor can measure PA levels at baseline, follow uptake through endogenous or overexpressed transporters upon addition to media, detect PA levels in diverse subcellular compartments in real-time, and monitor the effects of small molecule compound libraries on PA uptake and metabolism. These genetically encoded sensors will be broadly useful for studying the basic biology of PA metabolic and signaling pathways, as well as applied studies like pharmaceutical and genetic screens.

Our sensors are most sensitive to spermidine, spermine, and N^1^-acetylspermine (in that order), and to a lesser extent putrescine, agmatine, and cadaverine. They do not respond to ornithine. *A priori*, one should interpret iPASnFR signal rises as reflecting increased levels of the most responsive molecules, and signal dips as reflecting decreased levels. The normalization to fused mScarlet3 fluorescence helps control for differences in expression level and motion artifacts. In some circumstances, though, such as addition of excess putrescine, the sensor will primarily be exposed to this ligand, and fluorescence will primarily correspond to putrescine (albeit much less sensitively, given the sensor’s weaker response). While preparing this manuscript, a genetically encoded biosensor based on PotF was reported for agmatine detection^79^, displaying weak affinity and low signal change towards PAs. However, the authors only report and advocate its use for detecting agmatine.

We validated the iPASnFR-L sensors in ATP13A3-expressing cells by comparing side-by-side the uptake of green fluorescent PA probes and response of the sensor to the uptake of unlabeled PAs. With both methods we observed a dose-dependent increase in PA uptake that is ATP13A3-transport dependent, and confirmed that the sensor in cells selectively detects spermine and spermidine, but not putrescine. However, the window between ATP13A3-stable-OE and ATP13A3^D498N^-stable-OE observed with the iPASnFR-L in response to unlabeled PAs appears smaller than with the uptake of fluorescent PA probes. This can be easily explained by the differences between both methods. With iPASnFR-L we selectively monitor cytosolic increases in the free (unbound) PA levels, whereas with the uptake of fluorescent PA probes we determine the total cellular uptake, *i.e.*, encompassing free and bound PA probes present in endosomes, cytosol, and other compartments. Since iPAS-nFR-L exclusively responds to free cytosolic PAs, we have now unambiguously demonstrated for the first time that ATP13A3 transport activity elevates free cytosolic PA levels. Additionally, we have successfully targeted the sensors to subcellular compartments including the nucleus and mitochondria, providing the first proof-of-concept of organelle-specific free PA measurements. Further development of iPASnFR fusion constructs targeting a wider variety of organelles (*e.g.,* endoplasmic reticulum, Golgi apparatus, peroxisomes, early/late endosomes, and lysosomes) will provide deeper insights into how PAs enter, leave, and distribute within the cell.

To demonstrate a practical use of iPASnFR, we screened a compound library in search of metabolic signaling pathways potentially modulating downstream PA metabolism. Consequently, we identified multiple compounds that either increased or decreased cytosolic PA levels, revealing both known and unknown signaling pathways. A number of PA-decreasing compounds screened here converged in the PI3K/AKT/mTOR pathway and were found to significantly affect gene expression of PA-modulating genes in various cancer cell lines^80–92^. PA-synthesizing genes (*ARG1, ARG2, ODC1, SRM, SMS, AMD1*) were largely decreased in expression by these compounds, with CC-115, Apitolisib, and Torkinib decreasing expression of all PA-synthesizing genes in the pathway. Expression of *ODC1*, a rate-limiting enzyme in this pathway, was decreased in 17/22 (77%) of the experiments where differential expression for this gene was detected. Expression of at least 1 out of 4 PA-catabolizing genes was also increased by these compounds, with CC-115, CBL0137, and Omipalisib increasing expression of all catabolic genes examined, further depleting cellular PA contents. While the pattern for transporters was more complex, *SLC3A2* was most consistently decreased across experiments (22/28 = 79%), followed by *SLC18B1* (13/17 = 76%), *ATP13A2* (14/19 = 74%), and *ATP13A3* (15/21 = 71%), consistent with a decrease in PA uptake into the cytosol. However, in the case of CC-115, in studies where differential expression was detected, *ATP13A4* expression was increased in all cell types, *ATP13A3* was increased in the majority of cell types, but *SLC3A2*, *SLC18B1*, and *ATP13A2* show cell type-dependent up- or down-regulation, suggesting complex cell type-specific regulation of PA transport across different B-cell lymphoma cell lines. iPASnFR will be helpful in resolving the spatiotemporal changes induced by these drugs, including by what mechanism PAs are decreased, whether from efflux out of the cell or increased storage in intracellular organelles. These types of experiments could help resolve the seemingly complex effects of these drugs on transporter function and expression, to uncover the overall impact of these changes on PA levels and how this relates to therapeutic responses. In addition, given the complex architecture of tumors, application of iPASnFR *in vivo* will be critical in identifying cellular states and cell types within the tumor microenvironment that contribute to regulation of PA levels and how they might coordinate with one another.

There are numerous remaining mysteries of PA cell biology that the sensors can help solve. PAs have been implicated in regulation of the circadian clock in diverse organisms^93^, likely through a mechanism of reversible post-translational modification of clock proteins, as well as modulation of translational efficiency of their encoding mRNAs through ribosome shunting by disrupting interactions between the ribosome and 5’-UTRs^94^. There are many open questions, for instance: What are the mechanisms of the strong PA association with healthy aging^95^? What are the general effects of PAs on protein folding and stability, and how is this regulated in specific (and changing) circumstances^96^? Given the incredibly strong association of PA levels with cancer, why have attempts to drug this pathway failed^97^? How widespread are post-translational and post-transcriptional modifications of proteins and RNA by PA^10,98^? How closely synchronized are levels of distinct species of PAs among different organelles, and through what mechanisms^99^? Given the numerous roles of PAs in cell biology, how possible will drugging these pathways be^100^? What is the role of PAs in neuron-astrocyte signaling^101,102^? Many other questions remain^2^; in many ways, the rigorous study of PAs is in its infancy. Our sensors can be used to directly probe these mechanisms and open questions.

To answer these questions, future improvements to the iPASnFR family will include creation of i) higher-sensitivity and tailored-affinity variants, considering the relatively small dynamic range we saw with iPASnFR in mitochondria, ii) sensors in other color channels, iii) variants targeted to other organelles (endolysosomes, cell surface, *etc*.), iv) engineered sensors responding in different imaging modalities such as fluorescence lifetime^103^, and v) variants specific to spermine, spermidine, or putrescine, and discrimination of PA derivatives, such as acetylated PAs, which mediate a host of downstream pathways and also modulate PA biosynthesis itself^104,105^. Due to the broad selectivity of the currently available iPASnFR sensor to various PAs, albeit with different affinities, it remains challenging to convert the fluorescent signals into absolute PA concentrations. More selective PA sensors generated by mutagenesis or selection of other bacterial PA-binding proteins would circumvent this problem and provide PA-specific responses. Overall, this latest addition of iPASnFR to the current PA biology toolkit will greatly advance our understanding of polyamine functions on multiple physiological and pathological levels.

## DATA AVAILABILITY

All raw data are available upon request. Sensor sequences have been deposited in Genbank (iPASnFR-H: PQ241320, iPASnFR-L: PQ241321).

## CODE AVAILABILITY

The image analysis scripts are available on the lab’s GitHub page (https://github.com/LorenLoogerUCSD).

## REAGENT AVAILABILITY

Plasmids encoding iPASnFR-H or iPASnFR-L under several promoters and fusion proteins have been deposited at Addgene (#225235-225244).

## ACKNOWLEDGEMENTS

This work was supported by the Howard Hughes Medical Institute and startup funding from UCSD, as well research grants of the Fonds voor Wetenschappelijk Onderzoek (FWO) - Flanders (G009324N and G011424N) and the KU Leuven research grants C3/22/048 and C14/21/095 to PV. We would like to thank Chris van den Haute from the KU Leuven Viral Vector Core for generating stable cell lines. The Sanford Burnham Prebys Cancer Metabolism Core is supported by NCI Cancer Center Support Grant P30 CA030199. DAS is supported by NCI Research Specialist award R50 CA283813. We would like to thank Dr. Ilya Kolb for providing technical support on microscopy experiments.

## AUTHOR CONTRIBUTIONS

R.T., P.V., and L.L.L. designed the project. R.T. led the project. R.T. and L.L.L. designed iPASnFR-H/L. R.T. purified and characterized iPASnFR-H/L. R.T. made mammalian expression constructs for iPASnFR-H/L and conducted drug perturbation experiments in cultured cells. D.A.S. performed metabolomics. J.C. and M.D.J. performed cellular PA uptake assays. R.T. performed live imaging. R.T. conducted compound screens. R.T. and J.F.M. analyzed the results from the compound screen. J.F.M. conducted transcriptome analyses. R.T., J.F.M., P.V., and L.L.L. wrote the paper, with contributions from all the authors.

## METHODS

### Initial iPASnFR design

PcPotD is numbered such that the asparagine immediately following the leader sequence (KKLTIFFAAVFGAALLLYYVSLALNRTEGYVGK) is residue #2 (**Supp. Fig. 1**). cpsfVenus from iGluSnFR3^62^ was inserted between E284 and D285, flanked by the N-terminal linker (-LV-) and the C-terminal linker (-NP-), yielding PcPotD-cpsfVenus-E284. Other insertion variants shown in **Supp. Fig. 2** were designed similarly. PcPotD-cpsfVenus-E284 was synthesized (Integrated DNA Technologies) and cloned into the pBAD/His-B vector (Invitrogen) using Gibson assembly.

### Sensor optimization

Libraries of linker variants were constructed by a set of forward and reverse primers, each containing two consecutive NNK degenerate codons to replace the original linkers. These two primers were used to PCR-amplify cpsfVenus, followed by gel extraction and purification. The resulting linker libraries were inserted between E284 and D285 of PcPotD using Gibson assembly, transformed into *E. coli* TOP10 cells (Invitrogen), and plated onto agar plates with 100 µg/mL carbenicillin and 0.2% arabinose. Fluorescent colonies were picked into deep 96-well plates (Abgene) with 0.8 mL of Luria Broth (LB) media supplemented with 100 µg/mL carbenicillin and 0.2% arabinose and grown by rapid shaking at 37 °C for 18 h. Plates were centrifuged, bacteria washed with TBS (50 mM Tris, 300 mM NaCl, pH 7.5) twice, and pellets frozen at −80 °C for cell lysis overnight. The pellets were thawed, 0.5 mL ice-cold TBS added to each well, and crude lysates centrifuged. 90 µL aliquots of clarified lysates were transferred to black 96-well plates (Corning), and fluorescence was measured with a Tecan Spark plate reader before and after spermidine supplementation (10 mM final). Fluorescence ratio between pre- and post-spermidine addition was computed, and clones with high fluorescence change were recovered and sequenced in full (Plasmidsaurus).

### *In vitro* characterization

To purify PcPotD-cpsfVenus insertion variants and iPASnFRH/L, the corresponding pBAD/His-B plasmids were transformed into TOP10 cells and subsequently grown in 200 mL LB media with 100 µg/mL carbenicillin at 37 °C. Protein expression was induced by adding arabinose (0.2% final) at the OD_600_ of ∼0.3 for 12 h. Bacterial cultures were centrifuged, pellets were resuspended in PBS with protease inhibitor cocktail (04693159001, Roche), followed by sonication and centrifugation to collect supernatant. Clarified cell lysates were loaded onto Ni-NTA resins (Fisher Scientific) pre-equilibrated in PBS with 10 mM imidazole. Subsequently, bound lysates were washed with PBS with 25 mM imidazole and eluted with PBS with 250 mM imidazole. Fluorescent fractions were pooled, and buffers were exchanged into TBS using Amicon centrifugal filters (30 kDa cutoff, EMD Millipore).

To characterize the fluorescence response of purified proteins, increasing concentrations of polyamines (ornithine, cadaverine, agmatine, putrescine, spermidine, spermine, and N^1^-acetylspermine) and polyamine analogs (DFMO, AMXT-1501, Sardomozide, MGBG, Berenil, DENSPM, and MDL-72527) were prepared in TBS. Fluorescence of purified proteins (200 nM final) was measured using a Tecan Spark plate reader (excitation: 510 nm; emission: 540 nm at 5 nm intervals; 5 nm bandwidth) before and after compound addition with three technical replicates. To compute affinity, response curves were fitted to the Hill equation *Y* = *Y_max_* * *X* ^ *h* / *K_d_* ^ *h* + *X* ^ *h*, where *Y* is ΔF/F_0_, *Y_max_* is (ΔF/F_0_)_max_, *K_d_* is equilibrium dissociation constant (affinity metric), and *h* is the Hill slope. Excitation and emission spectra of purified proteins were also performed using the Fluorescence Intensity Scan function of the Tecan Spark: emission was scanned with 490 nm excitation and excitation with 540 nm emission (5 nm bandwidth) in the presence or absence of 10 mM spermidine. pH sensitivity of iPASnFR-H/L was measured in triplicate using purified sensors (200 nM final) in pH titration buffers (30 mM trisodium citrate, 30 mM borate) ranging from pH 4.0 to 9.0 (adjusted by either HCl or NaOH) in the presence or absence of 10 mM spermidine. p*K_a_* was calculated by fitting the pH titration curves to the sigmoidal function *Y* = *Y_max_* / 1 + 10 ^ (p*K_a_ –* pH), where *Y* is ΔF/F_0_, *Y_max_* is (ΔF/F_0_)_max_.

### Mammalian expression of iPASnFR-H/L

In general, HEK293 and U2OS cells (ATCC) were cultured in Opti-MEM (Gibco) supplemented with 5% fetal bovine serum (FBS, MP Biomedicals) and antibiotics (SV30079.01, Cytiva) at 37 °C with 5% CO_2_ in a humidified incubator. To cytosolically express iPASnFR-H/L in mammalian cells, their coding sequences were subcloned into pcDNA3.1 (Invitrogen) using Gibson assembly. To better quantify and compare their basal cellular brightness, mScarlet3 was C-terminally fused to iPASnFR-H/L, normalizing for expression in individual cells (pcDNA3.1-Cyto-iPASnFR-H/L-mScarlet3). Cells were transfected using TurboFect (Fisher Scientific) (as an example, DNA-TurboFect complex was prepared from 2 µg DNA and 6 µL TurboFect for 6-well transfection) following the manufacturer’s protocol. 24 h post-transfection, cells were trypsinized, rinsed with culture media, seeded onto clear bottom 96-well plates (Corning), and grown overnight. To assess the effects of PA-modifying compounds on endogenous PA concentrations, cells expressing iPASnFR-L were pre-treated with DFMO (1 mM - 10 mM) and AMXT-1501 (10 nM - 100 nM) either alone or in combination for 48 h before analysis.

### Imaging

Confocal images of cells (either untreated or treated) were acquired on a Keyence BZ-X810 microscope with 10x magnification 0.3 NA objective, the 49002 ET-EGFP fluorescence filter set (excitation: 470 nm/40 nm; dichroic mirror: 495 nm; emission: 525 nm/50 nm), and the 49031 ET-Cy3.5/Alexa568 filter set (excitation: 569 nm/25 nm; dichroic mirror: 588 nm; emission: 615 nm/40 nm) (Chroma). To account for regional variation, 3 regions were randomly selected in each well. For image analysis, a customized, automated pipeline was assembled using CellProfiler3.0^106^. Briefly, cells were segmented with the Robust Background algorithm, and the identified regions were masked in the green (525 nm emission) and red (615 nm emission) channels. The resulting images were used to compute the average background fluorescence in each channel, which was then subtracted from the original images. Lastly, green-to-red ratios were calculated for individual identified regions each corresponding to a single cell. Up to 500 data points (cells) were obtained and used to calculate mean green-to-red ratios per well. The image analysis pipeline is available on the lab’s GitHub page (https://github.com/LorenLoogerUCSD).

To visualize real-time PA uptake, HEK293 cells were transfected with pcDNA3.1-Cyto-iPASnFR-L-mScarlet3 in the presence or absence of plasmids encoding hATP13A2 or ATP13A3. To minimize cytotoxicity from excessive PA uptake, plasmids for iPASnFR-L and transporters were transfected at a 20:1 ratio. 24 h post-transfection, cells were harvested by trypsinization, seeded onto a 35 mm glass bottom dish (No. 1.0, MatTek) pre-treated with 50 µg/mL poly-D-lysine (Gibco) in PBS, and grown overnight. Live-cell imaging was performed on an inverted Olympus IX-83 microscope: Fluorescence was excited with a 470/24 nm LED (Spectra X Light Engine, Lumencor) through a standard ET-EGFP filter cube (excitation: 470/40 nm; dichroic: 495 nm; emission: 525/50 nm) and imaged with a 40x magnification 0.95 NA objective (Olympus) onto a scientific Complementary Metal-Oxide-Semiconductor (sCMOS) camera (ORCA Fusion, Hamamatsu) at 20 second intervals. Cells were first equilibrated for 30 min and imaged for 45 min in Opti-MEM (no pH indicator). Spermidine was added to a final concentration of 10 µM. Imaging was performed at room temperature (25 °C). Acquisition was performed using MetaMorph Advanced (Molecular Devices). Imaging analysis was conducted using ImageJ^107^. iPASnFR-L signal was quantified by subtracting background and manually selecting ROIs around individual cells.

### Metabolomics analysis

Polar metabolites extracted from cells and standards were prepared for Gas Chromatography-Mass Spectrometry (GC-MS) analysis as described^108^, with the addition of spermidine and spermine at the same concentration as putrescine. For GC-MS, a Thermo TG-S5SILMS column (30 m x 0.25 i.d. x 0.25 μm) installed in a Thermo Scientific TSQ 9610 GC-MS/MS was used. The GC was programmed with an injection temperature of 300°C and a 1.0 µL splitless injection. The GC oven temperature was initially 140 °C for 3 min, rising to 268 °C at 6 °C/min, and to 310 °C at 60 °C/min with a hold at the final temperature for 2 min. GC flow rate with helium carrier gas was 60 cm/s. The GC-MS interface temperature was 300 °C and (electron impact) ion source temperature was 200 °C, with 70 eV ionization voltage. Standards were run in parallel with samples. Metabolites in samples and standards were detected by MS/MS using precursor and product ion masses, and collision energies shown in the attached table (**Supp. Table 1**). Sample metabolites were quantified using calibration curves made from the peak areas of standards in Thermo Chromeleon software. Data were further processed to adjust for the relative quantities of metabolites in the standards, and for recovery of the internal standard (L-norvaline).

### Cellular polyamine uptake assay

HEK293T stably expressing wild-type ATP13A3 or catalytically dead variant D498N were generated by Leuven Viral Vector Core, *via* lentiviral transduction (Addgene plasmid ID for wild-type ATP13A3: #195849; ATP13A3-D498N: #195850). HEK293T cells were cultured in Dulbecco’s Modified Eagle Medium (DMEM) high glucose culture medium (Gibco), supplemented with 2% Penicillin-Streptomycin (Merck), 2 µg/mL puromycin (Invivogen) and 8% heat-inactivated FBS, south America origin (PAN BioTech). To perform transient transfection, culture medium was replaced by that without puromycin. The ATP13A3-stable-OE and ATP13A3^D498N^-stable-OE cells were transiently transfected with pcDNA3.1-Cyto-iPASnFR-L-mScarlet3, using TurboFect (Fisher Scientific) at 1:3 ratio in 15 cm cell culture dishes (Greiner) following the manufacturer’s protocol. 48h after transfection, cells were trypsinized and seeded in 12-well plates (Greiner) to reach 70% confluency the next day. To perform polyamine uptake, cells were treated with 1 mM aminoguanidine (Merck) with or without 100 nM AMXT-1501 (kindly provided by Aminex Therapeutics) 1h prior to addition of either unmodified or BODIPY-labeled PAs (Merck) at 37 °C and 5% CO_2_. After 0.5 h uptake, cells were trypsinized with TrypLE (Gibco), washed with PBS and resuspended in PBS supplemented with 0.1% Albumin Fraction V. (Carl Roth) and 2 mM EDTA (Calbiochem). Cell suspensions were filtered through a nylon filter to remove clumps. The samples were kept on ice until acquisition using Cytek Aurora Spectral Flow Cytometer (Cytek Biosciences). Mean fluorescent intensities of 10000 events per sample were recorded. Data analysis was performed with SpectroFlo (Cytek Biosciences).

### Small compound screen

To identify molecular interplays between major signaling pathways and intracellular PA metabolism, a library of 1,268 compounds (L2130, Anti-Cancer Metabolism Compound Library, TargetMol) was used. HEK293 cells were transfected with pcDNA3.1-Cyto-iPASnFR-L-mScarlet3 and seeded onto 96-well plates as described earlier. 48 h post-transfection, compounds were applied to cells at 2 µM (0.01% DMSO final). Images were taken at three time points (24, 48, 72 h post-treatment) on a Keyence microscope using a 10x objective, and the iPASnFR-L signal was processed as described above. Each plate contained two wells of DMSO-treated cells, which were used as a reference to obtain fold change values for individual compounds. Three independent screens were performed as experimental replicates, and compounds with fold change significantly different from 1.0 were determined using single-sample *t*-tests with a threshold of *P* < 0.05. We first collected the top 25 compounds increasing PA levels as determined by iPASnFR-L signal, as well as the top 25 compounds decreasing it. We then assessed the enrichment of specific pathways or targets associated with these 2 compound sets by conducting chi-squared or Fisher’s exact tests on contingency tables containing counts reflecting the frequency of each pathway or target term, compared to the remaining compounds. Fisher’s exact test *P*-values were used when any expected values in the contingency table were less than 5. Comparisons were not made for pathways or targets not represented in the combined set of top 50 compounds. The schematic of **Fig. 3A** was created with BioRender. The top 25 PA-increasing and -decreasing compounds were further evaluated for effects on expression of polyamine-associated genes documented in the literature. On August 15, 2024, the Gene Expression Omnibus (GEO) datasets database was searched for each of the top 25 PA-decreasing or -increasing compounds^109,110^. Studies were restricted to high-throughput sequencing of human-derived cells or cell lines. GEO2R was used to perform differential expression testing with DESeq2^111^ at a false-discovery rate cutoff of 0.1. Data was included for different doses, treatment durations, or cell lines to assess consistency of drug effects on gene expression. This gene set included PA synthetic genes (*ARG1, ARG2, ODC1, SRM, SMS, AMD1*), catabolic genes (*SAT1, SAT2, PAOX, SMOX*), and transporters (*SLC3A2, SLC18B1, ATP13A2, ATP13A3, ATP13A4*). The drug effect on gene expression was recorded as “decreased” or “increased” if gene expression was reduced or increased by the drug, respectively.

### Statistics and reproducibility

All errors presented in the text are s.e.m. All titration experiments with purified proteins (**Fig. 1D and Supp. Figs. 2B-C, 3, 6B**) had 3 technical replicates. Quantification of basal iPASnFR-H/L signal in HEK293 cells was independently performed 2-3 times (**Fig. 2B, D**). For quantifying PA uptake by ATP13A3-stable-OE and ATP13A3^D498N^-stable-OE HEK293T cells, 3-4 independent experiments were performed (**Fig. 3A-B**). Live-tracking of iPASnFR-L in HEK293 cells was obtained from at least 2 independent experiments, and data points for individual ROIs were pooled per condition (**Figs. 3D and 4B**). Compound screens were independently performed 3 times (**Fig. 5**). Statistical analyses were performed in GraphPad Prism or R, and types of tests performed were described in the main text and legends. Data distribution was initially assessed for normality using the Shapiro-Wilk test. If normal, single-sample *t*-tests (for fold change from the value 1.0) or two-tailed unpaired *t*-tests (for pairwise analyses) were performed. If not normal, 2-tailed Mann–Whitney *U* tests were performed (for pairwise analyses). Statistical significance was represented as follows: * *P* < 0.05; ** *P* < 0.01; *** *P* < 0.001; **** *P* < 0.0001; non-significant (ns).

---

**Supplementary Figure 1:**
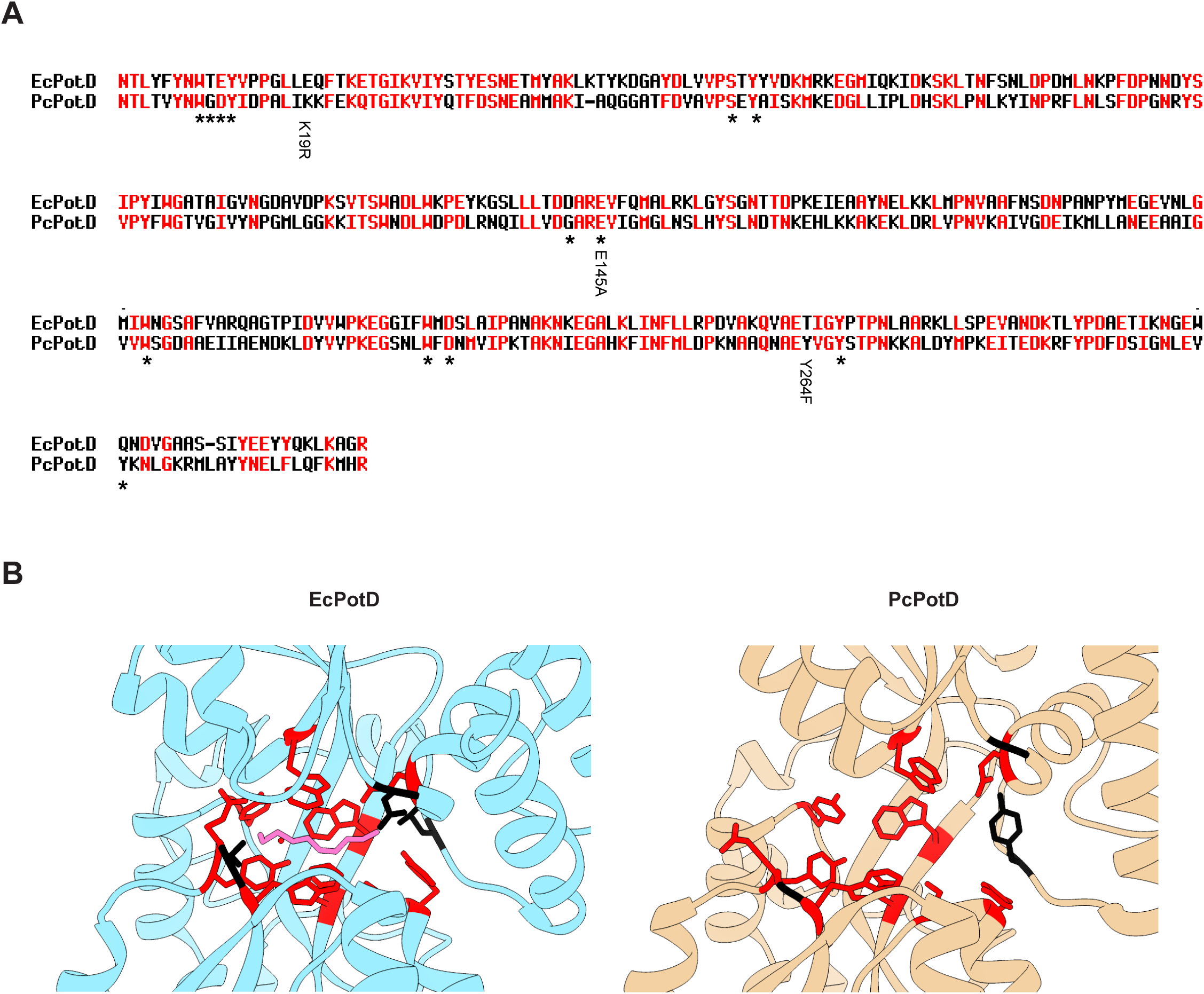
Primary sequence alignment of EcPotD and PcPotD. (**A**) Primary sequence alignment of EcPotD and PcPotD using MultAlin^112^. Amino acids that are either identical or similar are represented in red and those dissimilar in black. Amino acid residues involved in spermidine binding in EcPotD are indicated by asterisks. Mutations found in iPASnFR-H/L are indicated (related to **Fig. 1A**). (**B**) Left: crystal structure of EcPotD and spermidine in complex (PDB: 1POT); right: AlphaFold2-predicted structure of PcPotD binding pocket. Binding residues marked in (**A**) are represented in red and black as denoted in (**A**).

**Supplementary Figure 2:**
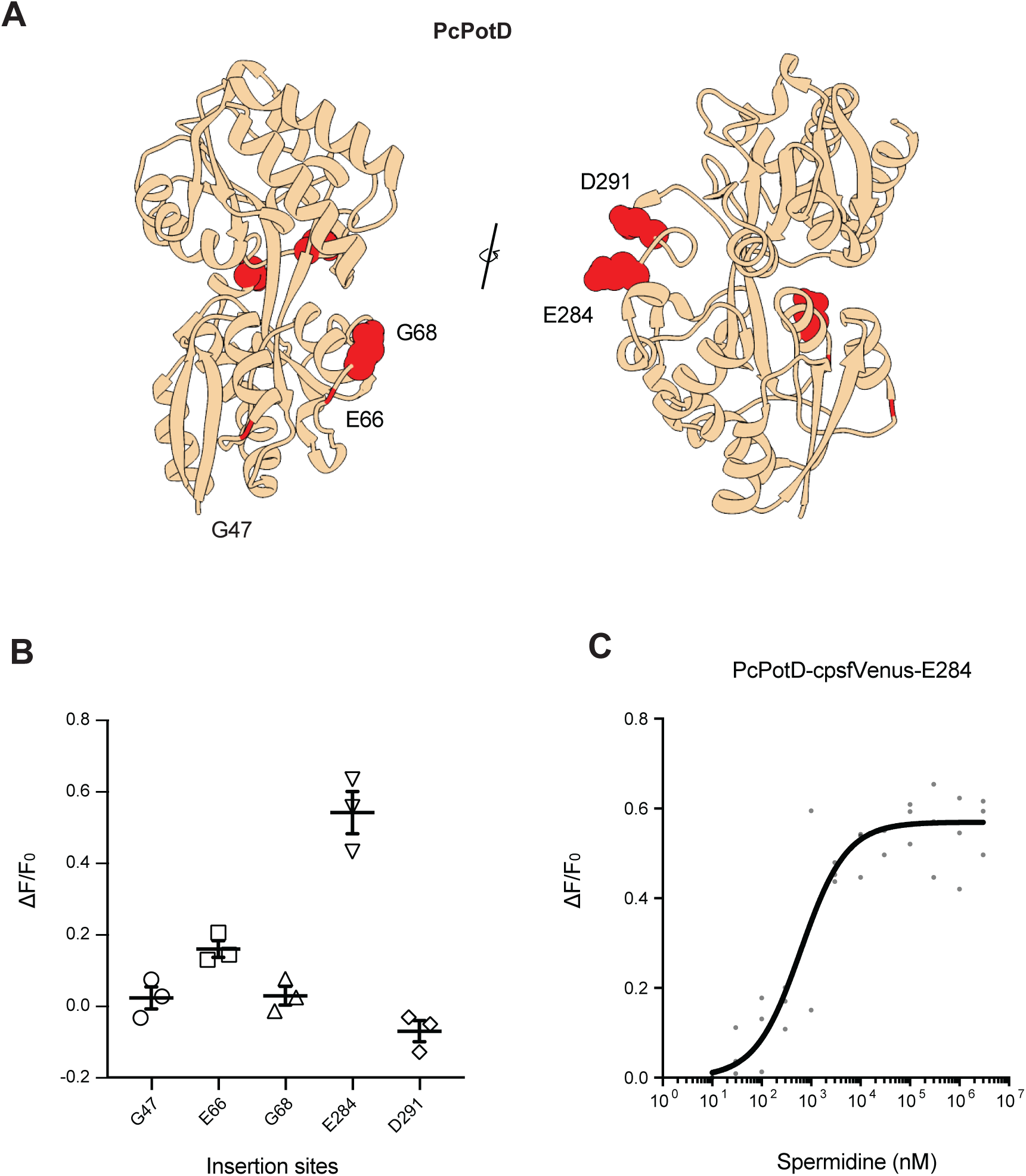
Fluorescence response of insertion variants of PcPotD-cpsfVenus. (**A**) AlphaFold2-predicted model of PcPotD. All insertion variants tested in this study (G47, E66, G68, E284, and D291) are shown in red. (**B**) Fluorescence increase (ΔF/F_0_) of purified prototype PcPotD-cpsfVenus insertion variants in the presence of 1 mM spermidine relative to its absence (*n* = 3). (**C**)Titration of purified PcPotD-cpsfVenus-E284 with spermidine (*n* = 3).

**Supplementary Figure 3:**
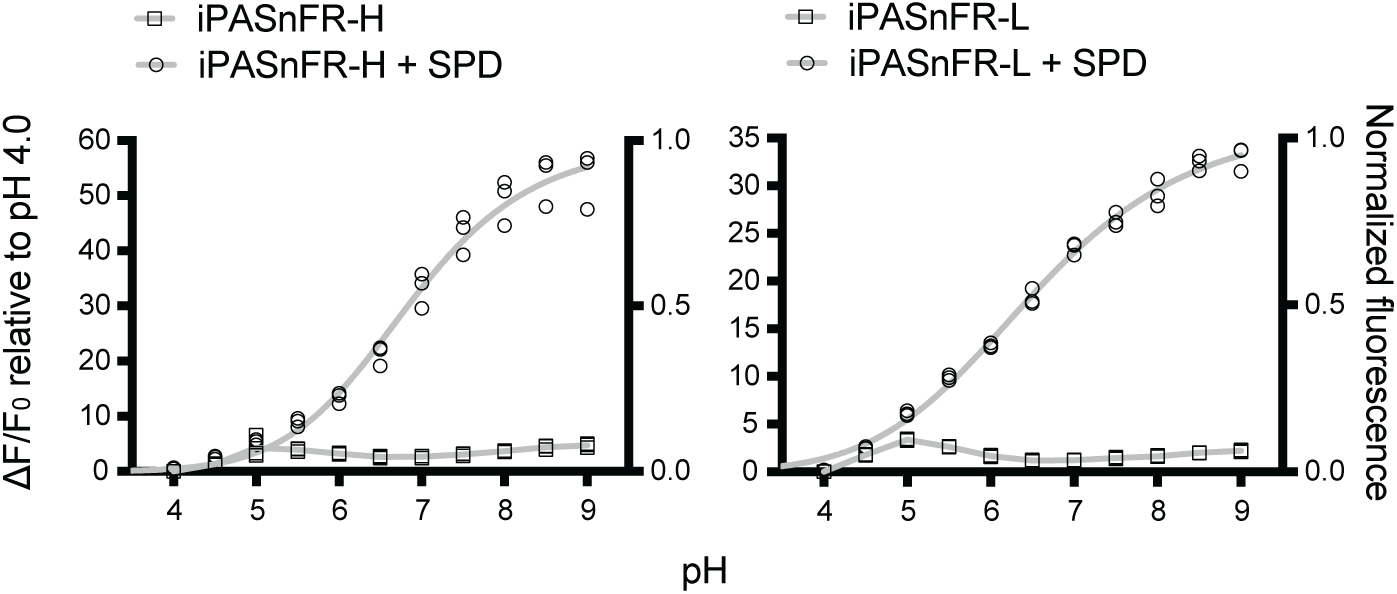
pH sensitivity of iPASnFR-H/L. iPASnFR-H/L fluorescence in the presence or absence of 10 mM spermidine, in buffers clamped at various pH values. Three independent titrations were performed, and both ΔF/F_0_ relative to pH 4.0 and fluorescence normalized to maximum response are shown.

**Supplementary Figure 4:**
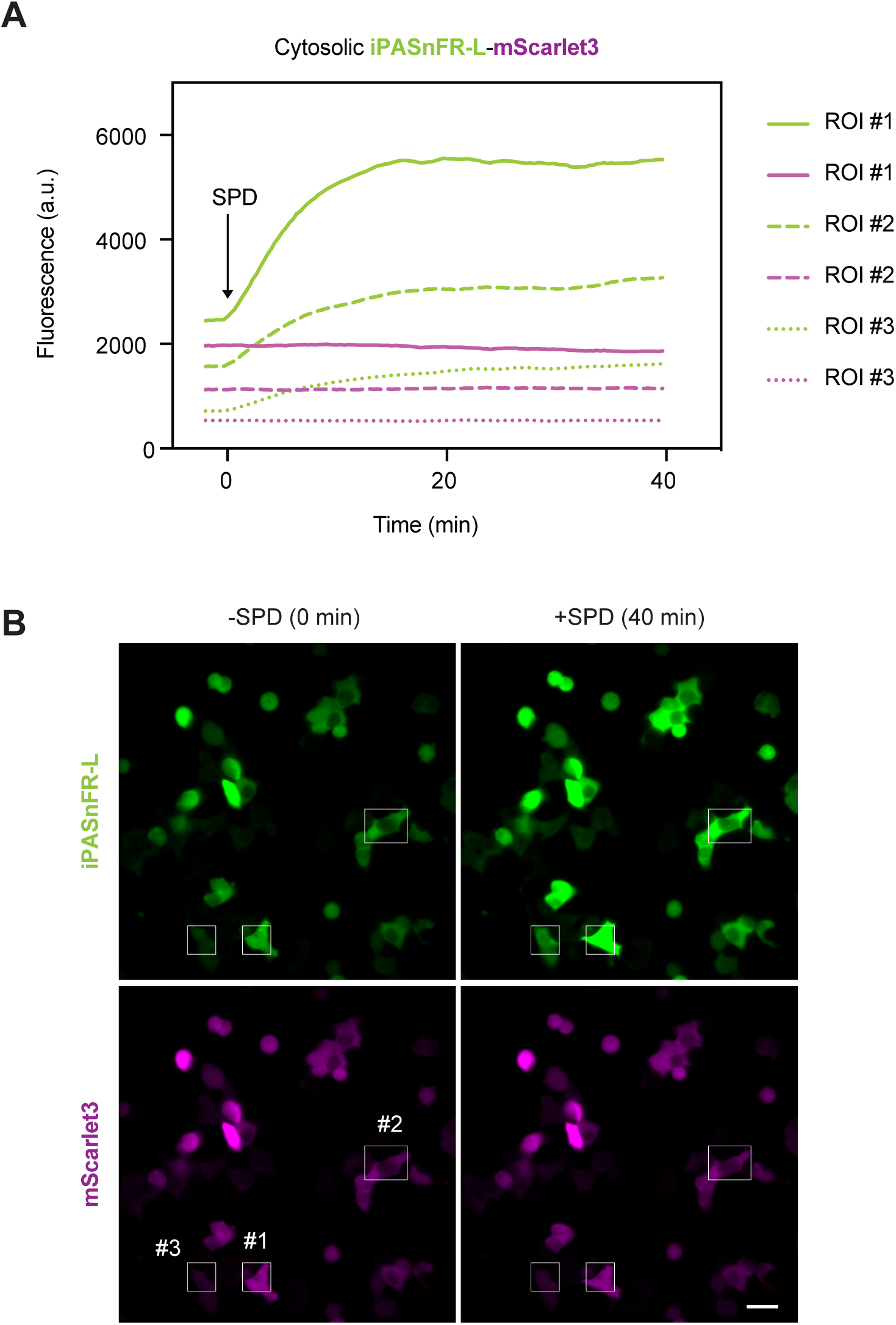
Example traces of iPASnFR-L-mScarlet3 in the cytosol upon spermidine addition. **(A)** Example raw traces of cytosolic iPASnFR-L-mScarlet3 signals in HEK293 cells expressing ATP13A3 upon spermidine addition (related to **Fig. 3C-D**). Three ROIs were individually plotted. Light green: iPASnFR-L emission; magenta: mScarlet emission. Matching line patterns indicate matching ROIs. (**B**) Raw image showing the corresponding ROIs traced in (**A**). Two time points (left: pre-spermidine addition; right: post-spermidine addition at 40 min) are shown. The same image was also used in **Fig. 3C**. Scale bar, 20 µm.

**Supplementary Figure 5:**
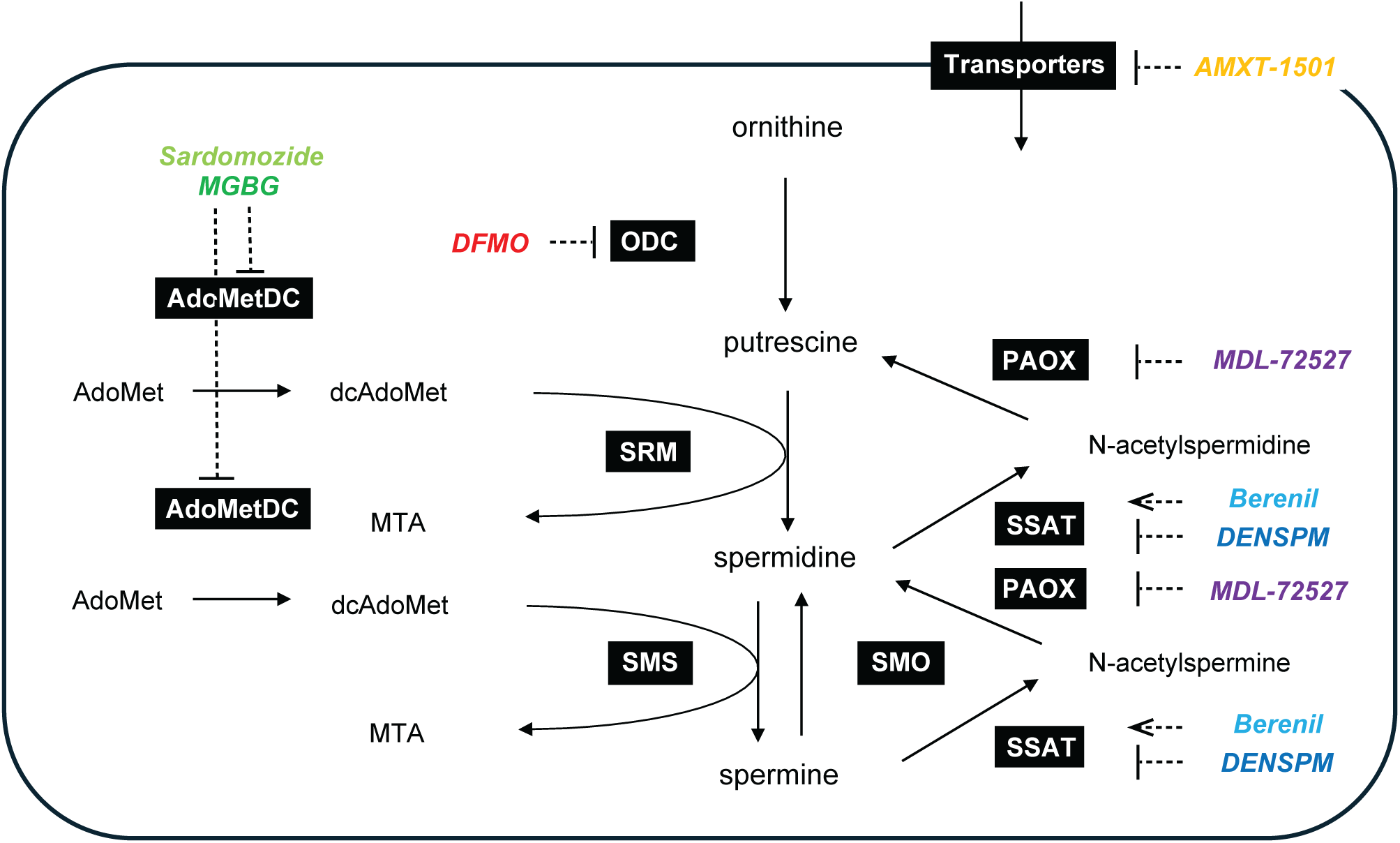
Targets of existing PA pathway-modifying compounds in PA metabolism. MGBG: methylglyoxal-bis(guanylhydrazone), a competitive inhibitor of AdoMetDC; PAOX: polyamine oxidase; SSAT: spermidine/spermine N^1^-acetyltransferase; SMO: spermine oxidase. The other acronyms are defined in the main text.

**Supplementary Figure 6:**
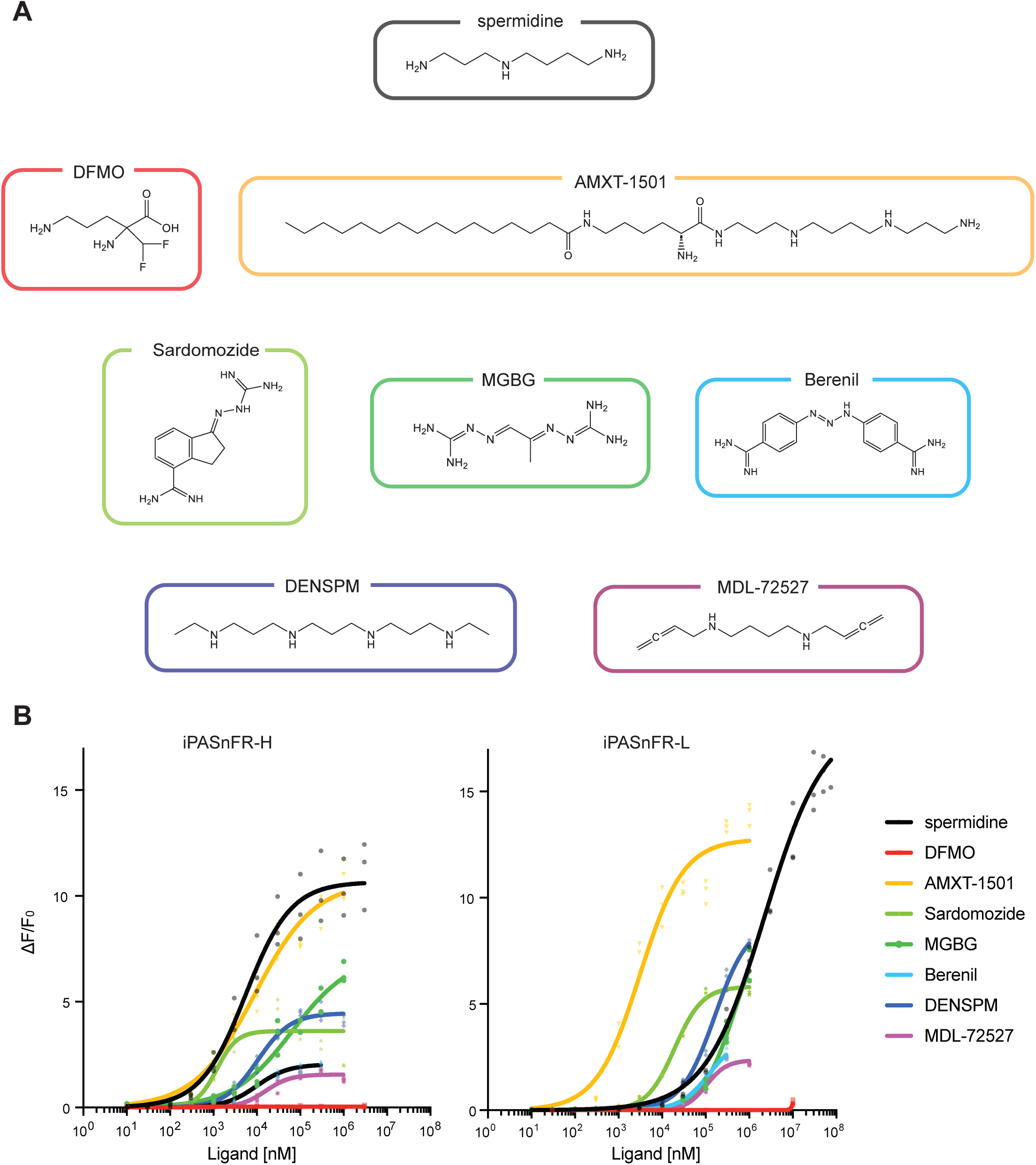
Fluorescence response of iPASnFR-H/L polyamine-modifying compounds. (**A**) Chemical structures of each drug shown in **Supp. Fig. 5**. Note that DFMO mimics ornithine, while others are analogs of spermidine and/or spermine. (**B**) *In vitro* titration of purified iPASnFR-H/L proteins with PA-modifying compounds shown in (**A**). Three independent titrations were performed for each compound, shown together.

**Supplementary Figure 7:**
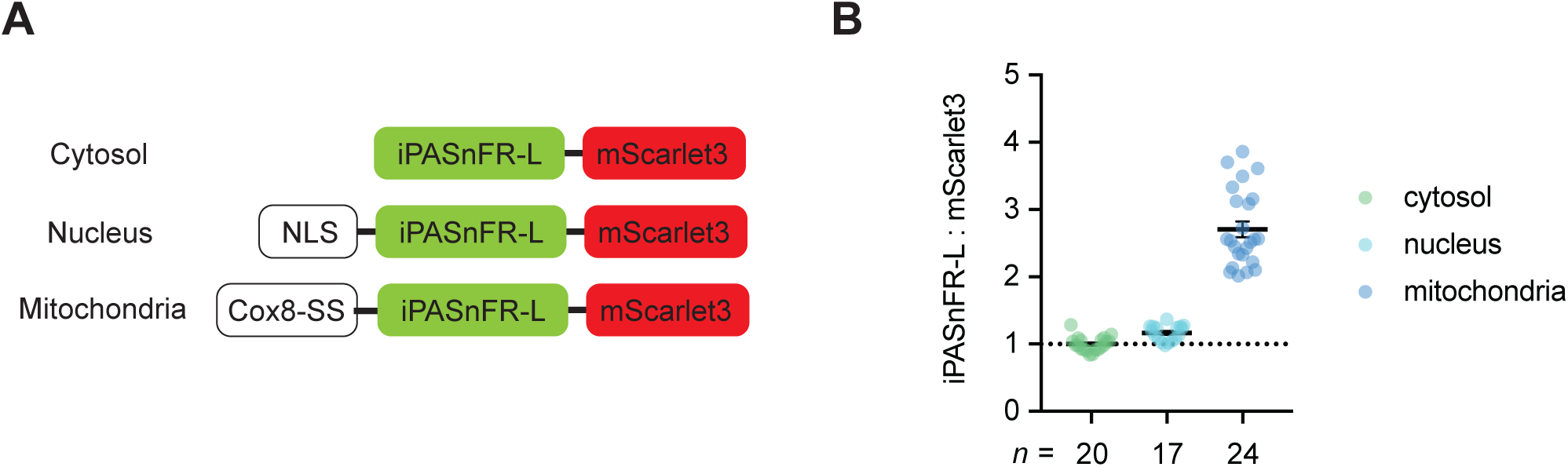
Relative subcellular concentration of polyamines. (**A**) Design of subcellularly targeted iPASnFR-L-mScarlet3 constructs. (**B**) Ratio of iPASnFR-L:mScarlet emission in the cytosol (*n* = 20), nucleus (*n* = 17), and mitochondria (*n* = 24). ROIs corresponding to single cells were collected from two independent experiments.

**Supplementary Figure 8:**
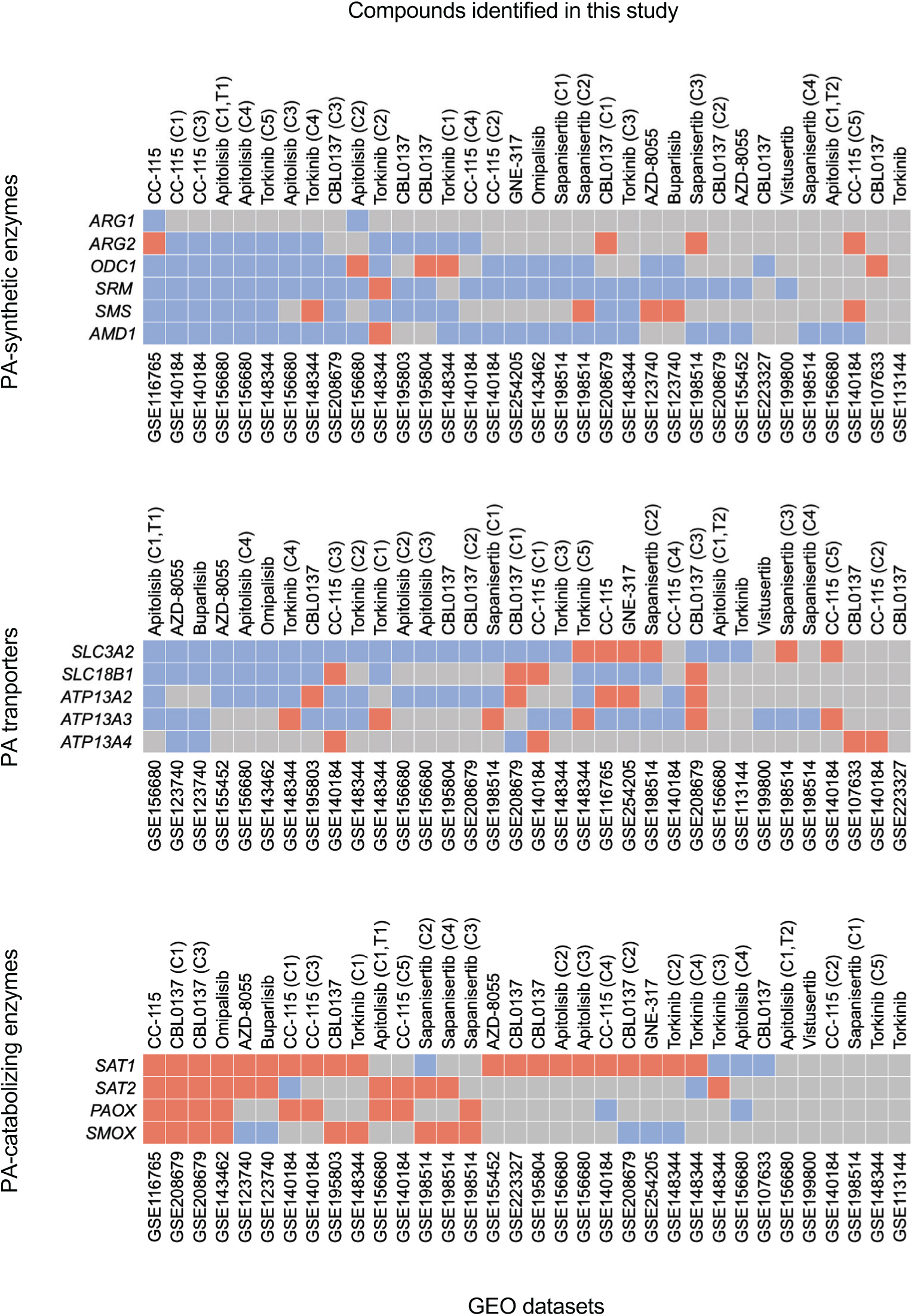
Effect of select top 25 polyamine-decreasing compounds on expression of polyamine-associated genes. Summary of PA-associated genes differentially expressed following treatment with the indicated compound from the top 25 PA-decreasing compounds. A search of the GEO database produced 16 studies consisting of 28 experiments with significant differential expression of these genes by treatment with CC-115, Apitolisib, Torkinib, CBL0137, GNE-317, Omipalisib, Sapanisertib, AZD-8055, Buparlisib, or Vistusertib^80–84,84,86–92,109,111^. The majority of cell types assessed were cancer cell lines, including those derived from gastrointestinal stromal tumors (GIST-T1, GIST-T1/670, TIST-T1/816), B-cell lymphoma (GRANTA-519, JEKO-1, Ramos, RPMI-8226, U266B1, SU-DHL-4), breast cancer (94T778, SKBR3, AU565, SUM52PE), and melanoma (A375, SKMEL28, Mel202). This included experiments in which treatments were given with the same drug in multiple cell types (CC-115 (GRANTA-519, JEKO-1, Ramos, RPMI-8226, U266B1), Apitolisib (GIST-T1, GIST882, GIST-T1/670, GIST-T1/816), Torkinib (94T778, A375, SKMEL28, SKBR3, AU565), CBL0137(SU-DHL-4, Raji Burkitt Lymphoma, JEKO-1, SKNMC, A673), Sapanisertib (ATRT-TYR molecular subgroup, ATRT-SHH, ATRT-MYC (CHLA06), ATRT-MYC (CHLA266))) or in the same cell types but with different treatment durations (Apitolisib (GIST, 4hr and 24hr)). Differential gene expression was determined with DESeq2^111^ to identify significantly altered PA genes (FDR < 0.1). Genes were grouped into sets based on role in PA biology, including PA synthetic genes (*ARG1*, *ARG2*, *ODC1*, *SRM*, *SMS*, *AMD1*) (Top panel), catabolic genes (*SAT1*, *SAT2, PAOX*, *SMOX*) (Middle panel), and transporters (*SLC3A2*, *SLC18B1*, *ATP13A2*, *ATP13A3*, *ATP13A4*) (Bottom panel). Drug effect is indicated by color, where blue indicates significantly decreased expression and red significantly increased expression. Studies are sorted separately for each gene set in descending order based on the total number of genes decreased (top and bottom panel) or increased (middle panel) within each gene set. In cases where multiple drug doses, incubation times, or cell types were tested in the same GEO record, these are differentiated by labels next to the drug name where “C” indicates cell type and “T” incubation time. These labels are followed by a number to indicate a different cell type or time point used for the same drug in the same study.

**Supplementary Table 1:**
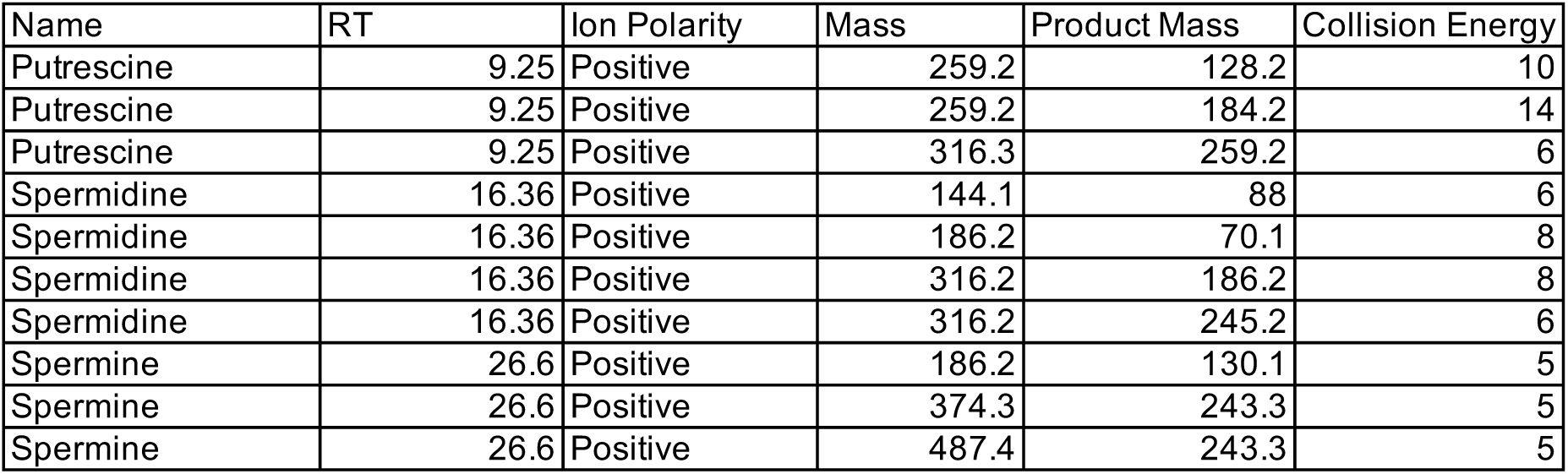
Precursor and product ion masses, and collision energies of putrescine, spermidine, spermine.

## Notes

### Competing Interest Statement

The authors have declared no competing interest.

### Summary of Updates

Author descriptions and figure labels were modified. GenBank accession numbers were updated.

